# Deep generative modeling reveals maturation-linked pairing signatures in human antibodies

**DOI:** 10.1101/2025.08.24.671684

**Authors:** Lea Bröonnimann, Thomas Lemmin, Chiara Rodella

## Abstract

Understanding how antibody heavy and light chains pair is critical for decoding immune repertoire architecture and designing therapeutic antibodies. However, most antibody sequence databases lack paired chain information. To address this gap, we developed a two-stage deep learning framework. First, we pre-trained separate transformer-based language models on large corpora of unpaired heavy and light chain sequences to capture patterns of gene usage and somatic hypermutation. These models were then integrated via lightweight adapters into a sequence-to-sequence model trained in a machine translation setting, enabling light chain generation conditioned on heavy chain input. Although native light chain recovery was moderate, the model consistently captured functionally meaningful constraints: generated sequences exhibited high germline identity, improved structural quality of predicted folds, and broader coverage of framework and CDR regions. Immunologically, heavy chains from memory B cells preferentially generated light chains with more restricted V gene usage, reflecting maturation-dependent selection. Additionally, generated ***κ*** light chains displayed a trimodal similarity distribution, suggesting distinct functional pairing modes ranging from promiscuous to highly specific. This work shows that sequence-to-sequence modeling can uncover inter-chain dependencies and generate structurally and immunologically plausible antibody pairs, providing a foundation for computational repertoire analysis and therapeutic design.

**Highlights:**

- A deep generative modeling framework enables conditional generation of light chains from heavy chains, leveraging unpaired data.
- Conditioning enhances the structural quality and germline coherence of predicted antibodies.
- Memory B cell-derived heavy chains preferentially generate light chains with restricted V gene usage, consistent with maturation-dependent selection.
- Generated *κ* light chains show a trimodal similarity distribution, suggesting discrete pairing modes ranging from promiscuous to highly specific.

## 1 Introduction

Antibodies are critical effector molecules of the adaptive immune system, produced by B cells to specifically recognize and neutralize pathogenic threats [1]. The canonical antibody structure consists of two identical heavy chains (HCs) and two identical light chains (LCs), arranged in a characteristic Y-shaped configuration that spatially separates antigen recognition from effector functions [1–3]. The immense diversity of antibodies arises from precise genetic mechanisms including V(D)J recombination, junctional diversity, and somatic hypermutation [3, 4]. HC construction involves the selection and joining of V, D, and J gene segments with random non-templated nucleotide additions at junctions, while LC rearrangement follows a similar but simpler VJ process that lacks D genes and involves fewer insertions and deletions [2, 5, 6]. Consequently, LCs are significantly less diverse than HCs, with mounting evidence suggesting that this constraint reflects the evolutionary pressure for LC rearrangements to minimize self-reactivity of B cell receptors (BCRs) [7, 8]. The variable domains of both HCs and LCs are organized into alternating framework regions (FRs) that maintain structural integrity and complementarity-determining regions (CDRs) that constitute the antigen-binding site, where the six CDRs collectively form the paratope responsible for antigen recognition [9, 10]. The remarkable combinatorial diversity of antibodies enables recognition of virtually any foreign epitope [3, 11]. Understanding how this diversity is organized and utilized within antibody repertoires is therefore essential for advancing fundamental immunology, rational vaccine design, and the development of engineered therapeutic antibodies with enhanced specificity and reduced immunogenicity [12, 13].

The advent of next-generation sequencing (NGS) has revolutionized the study of antibody repertoires, allowing high-throughput characterization of immune responses at an unprecedented scale [12]. By sequencing the variable regions of HCs and LCs, researchers can map the diversity and evolution of antibody responses in health and disease. However, NGS generates massive datasets that present significant analytical challenges. Although these technologies can efficiently capture the sequence information from millions of individual B cells, they typically lose the natural pairing information between HC and LC during the sequencing process [14–16]. This loss of pairing information creates a critical knowledge gap, as the function and specificity of an antibody critically depend on the correct association of its HCs and LCs [17, 18]. Consequently, computational immunology faces a fundamental challenge: how to accurately reconstruct or predict these essential pairing relationships from sequence data alone. The scale and complexity of antibody repertoire datasets further compound this challenge, necessitating advanced computational approaches that can effectively process and interpret vast amounts of immunological data [12, 19, 20].

Recent advances in deep learning, particularly the emergence of language models, have demonstrated remarkable capabilities in extracting meaningful patterns from large-scale biological datasets [19–21]. The application of these models to biological sequences is conceptually grounded in the principle that amino acid sequences encode protein structure and function through positional context and relationships, analogous to how words in sentences convey meaning through order and context [20, 22]. These models, inspired by natural language processing techniques, have been successfully applied to biological sequences, revealing hidden structures and relationships within repertoires. Although studies on protein and antibody language models have shown substantial progress [21, 23–28], there has been limited exploration into the specific task of predicting HC-LC pairings and generating one chain given the other [22, 29]. This gap is significant because certain HCs demonstrate strong preferences for specific LCs, suggesting that the pairing itself can be crucial for developing stable, efficacious therapeutic antibodies [30, 31]. It has been observed that many antibody pairs naturally favor cognate HC-LC pairing [31] and that randomly paired HC and LC often result in autoreactive or non-functional B cell receptors [17]. This preference leads to higher yields of correctly assembled BsIgG, reducing the need for extensive engineering of antibody fragments (Fabs) to achieve the desired pairing [31]. This simplifies the development process and can improve the therapeutic efficacy of bispecific antibodies, making targeted treatments more accessible and effective. HC and LC pairing influences antibody structure by affecting the paratope and interface of the variable heavy (VH) and light chain (VL) domains [32] and specific amino acid properties can influence HC-LC compatibility and binding affinity [33]. Additionally, recent investigations demonstrate that memory B cells exhibit a pronounced heavy-light interdependence [34]. In these cells, the usage of an identical HC V gene is closely associated with the preferential selection of a specific LC V gene, a pattern that is markedly less evident in naive B cells. This suggests that in memory B cells, the HC effectively predicts the optimal LC, highlighting the role of evolutionary pressures in shaping immune responses.

In this work, we leverage deep learning approaches to investigate HC-LC pairing. We develop and evaluate a set of specialized transformer models: HeavyBERTa for HC sequences, LightGPT for LC sequences, classification models for predicting B cell developmental states, and a Heavy2Light encoder-decoder architecture that translates HC sequences into corresponding LC sequences. These models enable us to infer pairing relationships and improve our understanding of repertoire architecture. Our findings provide a framework for applying deep learning to antibody sequence analysis, with implications for both fundamental immunology and therapeutic antibody discovery.

## 2 Results

Understanding HC-LC relationships in antibody repertoires requires models that can capture both individual chain characteristics and inter-chain dependencies. However, paired HC-LC sequences are vastly outnumbered by unpaired sequences, with public databases containing billions of unpaired sequences but only a few million paired examples [35]. While it is feasible to train a translation model directly on this paired data, the relatively limited size restricts the capacity of large models to generalize effectively and fully capture repertoire diversity [36]. To address this imbalance and leverage the full extent of available data, we adopted a two-stage modeling strategy. First, we developed two domain-specific language models: HeavyBERTa, based on a masked language modeling (MLM) architecture for HCs, and LightGPT, a causal language model for LCs. These models were pre-trained separately on more than 99 million HC and 22 million LC sequences from the Observed Antibody Space (OAS [35]) database. The sequences were restricted to those derived from unsorted B cells, memory B cells, and plasma B cells isolated from peripheral blood mononuclear cells (PBMCs). To eliminate potential bias from immune activation, we exclusively used data from healthy human donors with no documented disease or recent vaccination history (Figure 1A and B; Methods 4.2.1; Supplementary Table A1).

**Fig. 1.**
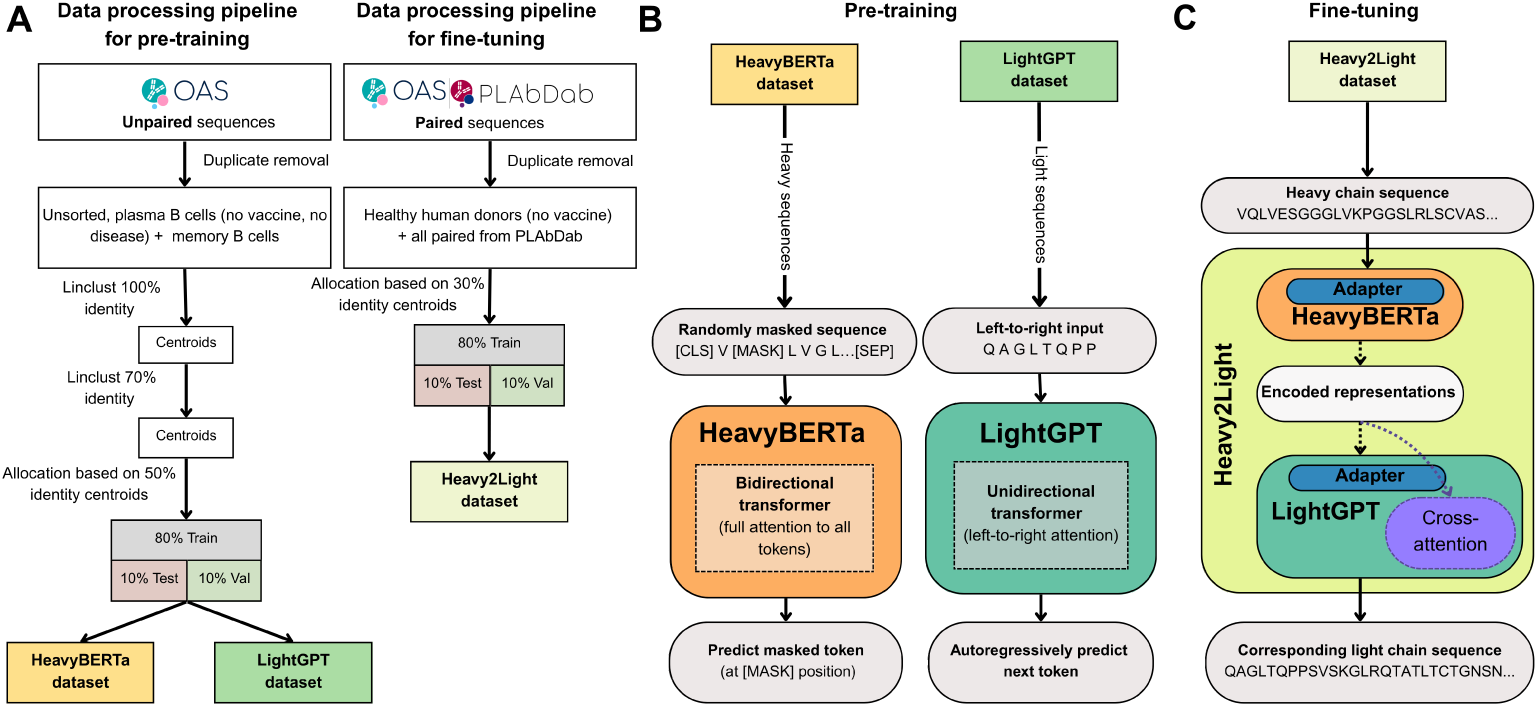
Data processing and training strategy. (A) Data processing pipeline showing the preparation of datasets for the pre-training and fine-tuning of our models. The CDR3 regions (CDRH3 for the HeavyBERTa dataset and CDRL3 for the LightGPT dataset) of unpaired sequences from the OAS (plasma B cells from unsorted samples, memory B cells) are clustered at 100%. Subsequently, the full sequence (heavy or light, for the respective dataset) of resulting centroids are clustered at 70% identity thresholds and allocated based on 50% centroids to create HeavyBERTa and LightGPT datasets. Paired sequences from OAS and PLAbDab are processed similarly with healthy human donor selection and 30% identity-based allocation, resulting in 80% training, 10% testing, and 10% validation splits for the Heavy2Light dataset. (B) Pre-training phase illustrating the training objectives for each model. HeavyBERTa employs bidirectional transformer architecture with masked language modeling on randomly masked HC sequences, while LightGPT uses unidirectional (left-to-right) transformer architecture for autoregressive next-token prediction on LC sequences. Both models learn sequence representations through their respective self-supervised objectives. (C) Fine-tuning phase demonstrating the encoder-decoder architecture where pre-trained HeavyBERTa (encoder) and LightGPT (decoder) are combined with cross-attention mechanisms and parameter-efficient adapter modules. Only the adapters and cross-attention parameters are trained, while the pre-trained model weights remain frozen. This Heavy2Light model translates input HC sequences into corresponding LC sequences, leveraging the learned representations from both pre-trained components.

HeavyBERTa was trained with two configurations to predict masked residues within HC sequences. Both variants demonstrated robust performance in this task (Supplementary Table A2). with the larger configuration achieving a marginally superior accuracy of (89.12%) compared to the smaller configuration’s (88.95%). The LightGPT model exhibited comparable performance in LC prediction, achieving an accuracy of 86.35%.

Our pretrained models demonstrate a strong ability to capture biologically meaningful features from antibody chain sequences. To examine the structure of these learned representations, we visualized the output embeddings using three complementary techniques: t-distributed Stochastic Neighbor Embedding (t-SNE), Principal Component Analysis (PCA), and Linear Discriminant Analysis (LDA). Across all visualization methods, both HeavyBERTa and LightGPT successfully clustered sequences according to their respective V gene families (Supplementary Figures A1, A2 and A3). Although the t-SNE and PCA projections were predominantly organized by V gene identity, we investigated whether the learned embeddings also captured information about J gene usage. Using LDA, we found clear separability of sequences based on their J gene segments (Supplementary Figure A3), with t-SNE and PCA visualizations further confirming these clustering patterns (Supplementary Figure A1 C, D and A2 C, D). These results demonstrate that our models encode immunogenetic information beyond V gene signatures, successfully capturing meaningful distinctions across different genomic loci.

Building on these findings, we examined whether the embeddings also reflected B cell developmental states [37]. B cell maturation is fundamental to understanding immune repertoires, as sequence differences between naive and memory B cells result from somatic hypermutation and selection during affinity maturation [38]. When we applied LDA to embeddings from both HCs and LCs, we found that both models could effectively distinguish between naive and memory B cell populations (Supplementary Figure A4). This separation indicates that the models have learned representations that capture biologically relevant differences related to B cell maturation states. This inherent ability to differentiate B cell states from sequence alone motivated us to finetune the HeavyBERTa and LightGPT models for direct classification of sequences according to their B cell origin. The HeavyBERTa model achieved high classification performance, with an overall accuracy of 92.31% and balanced performance across both naive (F1 score: 0.93) and memory (F1 score: 0.92) classes (Supplementary Table A3). This suggests the model effectively captures sequence features associated with somatic hypermutation patterns. In contrast, the LightGPT model demonstrated slightly lower performance (overall accuracy: 79.17%), with notably stronger identification of memory B cells (recall: 0.90) than naive B cells (recall: 0.58).

We further validated our classification results by examining HC-LC V gene pairing patterns. Memory B cells have been shown to exhibit a pronounced heavy-light interdependence, characterized by enhanced V gene coherence where identical HC V genes preferentially associate with specific LC V genes [34]. We tested whether this phenomenon would also be observed in previously unlabeled sequences when classified using our HeavyBERTa model. Our results confirmed the expected interdependence phenomenon in memory B cells (Supplementary Table A4, Methods 4.6.1). In the labeled OAS database, memory B cells demonstrated substantially higher V gene coherence (55.82%) compared to naive B cells (13.83%). This pattern was even more pronounced when applying our HeavyBERTa classifier to previously unlabeled sequences from unsorted B cell populations, where sequences classified as memory origin showed 80.57% coherence versus only 5.60% in those classified as naive origin. When combining all data, memory B cells maintained consistently higher coherence (60.63%) than naive B cells (11.90%). These findings validate both the biological phenomenon of enhanced HC-LC interdependence in memory B cells and demonstrate that our computational model successfully captures sequence features associated with B cell maturation states that correlate with established pairing preferences.

Given the demonstrated ability of our models to capture biologically meaningful patterns and the observed HC-LC interdependencies, we next explored whether these learned representations could be used for sequence generation tasks. To leverage underlying HC-LC pairing relationships, we developed Heavy2Light, an encoderdecoder architecture specifically designed to generate LCs conditioned on paired HC sequences, thereby explicitly modeling heavy-light interactions. Heavy2Light builds upon the strengths of our individually pre-trained HeavyBERTa and LightGPT models by integrating them into a paired architecture (Figure 1C). To learn the complex HC-LC pairing relationships, Heavy2Light was fine-tuned using a comprehensive dataset of paired antibody sequences. This dataset was compiled from the paired subset of the OAS [35], yielding 470,000 sequences from healthy human donors without vaccination or disease exposure, and the Patent and Literature Antibody Database (PLAbDab [39]), a self-updating repository containing over 150,000 paired antibody sequences curated from patents and academic literature (Methods 4.2.2, Supplementary Table A1). Fine-tuning was performed using adapter modules [40], which enable parameter-efficient training by introducing lightweight, trainable components into the frozen backbone of the model. This approach preserves the valuable pre-trained representations of HeavyBERTa and LightGPT while allowing the model to adapt to the specific pairing task. By updating only a small fraction of the model’s parameters, the use of adapters not only reduces computational requirements but also significantly accelerates training time compared to full model fine-tuning.

To assess the quality of our generated LCs, they were aligned with their reference LCs using global alignment (Methods 4.4). Sequence recovery between generated and true LC pairs shows considerable variation across antibody regions (Figure 2A). FR regions demonstrated moderate to high similarity, with FR2 achieving the highest mean identity of 72.90%, followed by FR3 at 70.08% and FR1 at 58.54%. The CDRs showed lower scores, with CDR2 at 40.29%, CDR3 at 33.40%, and CDR1 at 34.15%. Overall, the model achieved a total sequence identity of 60.73% between generated and true LC sequences (Supplementary Table A5). To uncover additional patterns and gain deeper insights into model behavior, we first classified the generated sequences as memory or naive using our maturation classifier, and subsequently divided them by LC isotype into *κ* and *λ* chains. The sequence identity distributions show distinct patterns across LC types and predicted maturation states. Both *κ* memory and naive B cell-predicted LCs exhibit a trimodal distribution, suggesting the presence of underlying structural or functional classes influencing pairing predictability (Figure 3B). Notably, the naive *κ* group shows a slight shift toward higher identity values, consistent with its higher mean similarity (67.05%) compared to the memory *κ* group (64.30%). In contrast, *λ* chains display a different behavior: Both *λ* groups lack clear multimodality and are dominated by a peak around 45%. Mean sequence identities were lower across both *λ* groups, with values of 54.91% for sequences predicted to originate from memory B cells and 54.42% for those from naive B cells. To investigate whether these patterns could be attributed to V gene family usage, we analyzed the distribution of *κ* V gene families across identity ranges. Although IGKV1 was overrepresented (Supplementary Figure A5), this alone did not explain the trimodal distribution, suggesting that more complex sequence–structure relationships underlie the observed trends. IGKV1 remained the most dominant V gene family across all generated LC sequences, followed by IGKV3, regardless of whether the heavy and LCs were predicted to have matching or mismatching maturation states. Specifically, IGKV1 accounted for 20.2% of sequences in the matching group and 11.9% in the mismatching group, while IGKV3 represented 15.7% and 12.2%, respectively (Supplementary Figure A6A). This pattern was consistent with the distribution observed in the true LCs, where IGKV1 was the most frequent gene family in both the test set (45.03%) and training set (30.58%). IGKV3 was the second most common, comprising 15.07% of the test set and 26.33% of the training set (Supplementary Figure A6 B and C).

**Fig. 2.**
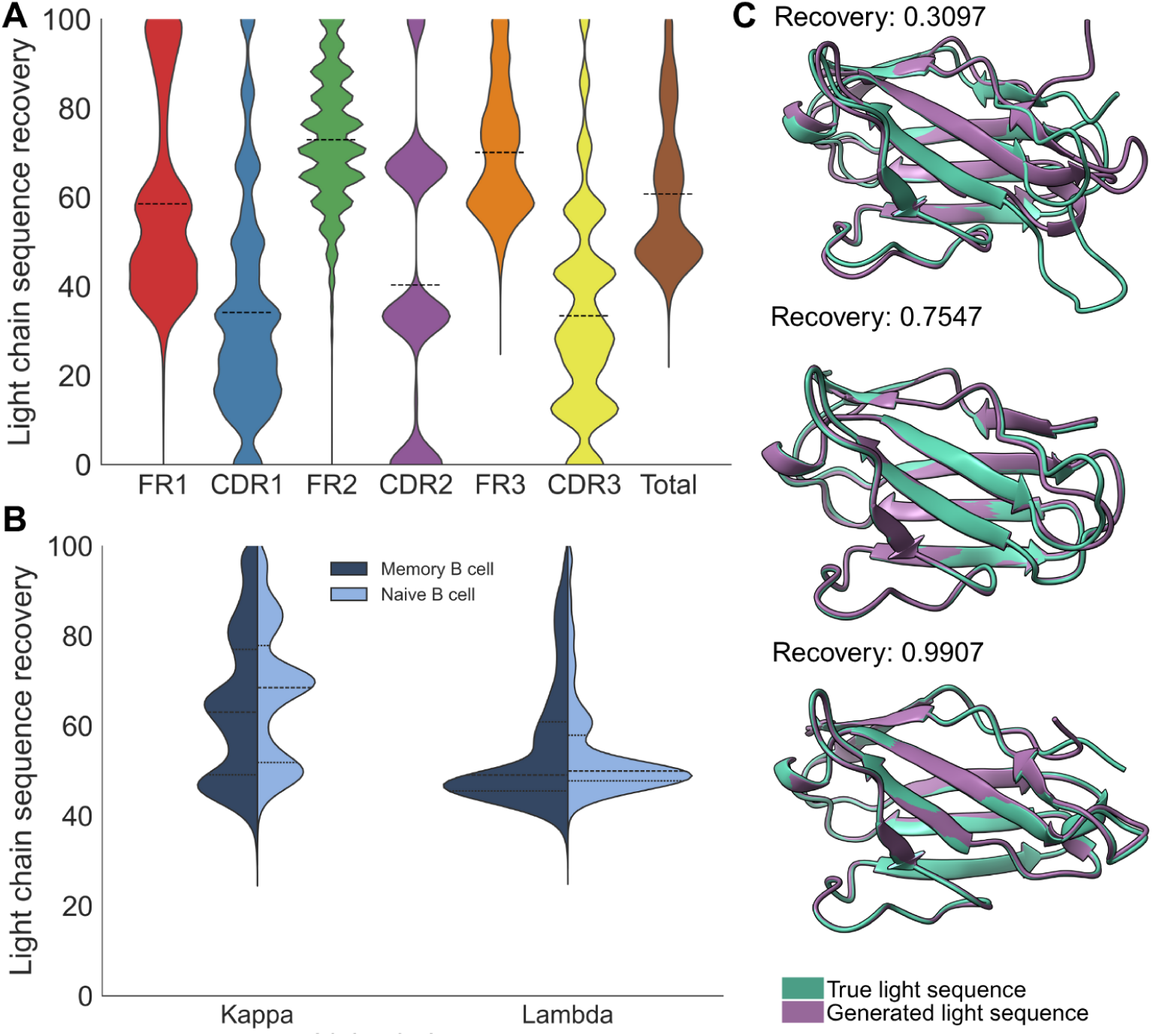
Characterization of conditionally generated LC sequences based on HC input. (A) Sequence identity (%) between generated LCs and their corresponding true counterparts across FR regions (FR1-3), CDRs (CDR1-3), and the complete variable domain. (B) Sequence identity distributions stratified by B cell maturation state (naive in dark blue and memory in light blue) and LC isotype (*κ* or *λ*). (C) Structural superposition of generated LCs (purple) with their true counterparts (teal) representing three distinct recovery levels. Sequence recovery is given next to each structure.

**Fig. 3.**
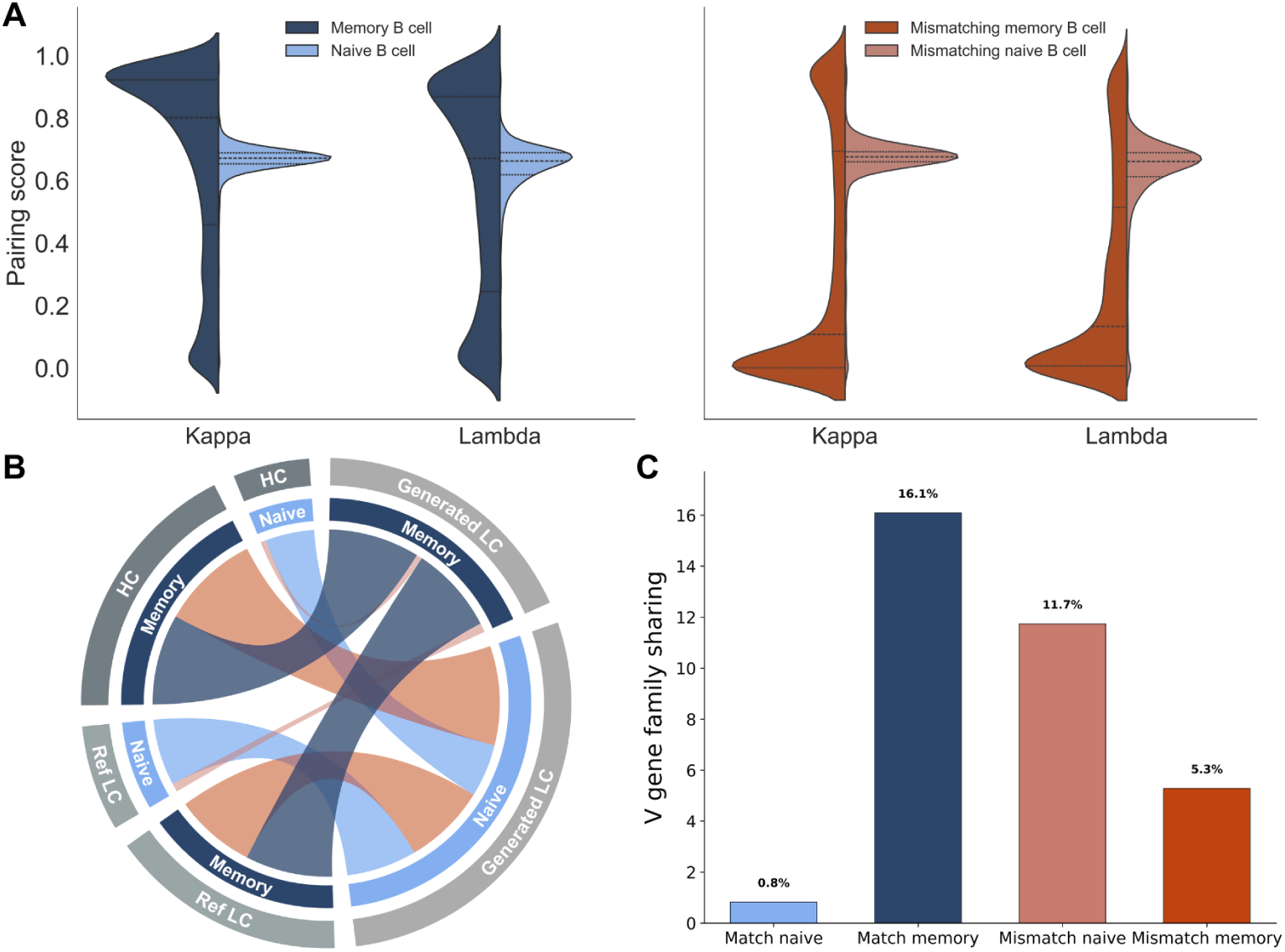
HC–LC pairing patterns of conditionally generated LC sequences. (A) Pairing scores between conditionally generated LC sequences and their corresponding HCs. (B) Chord plot illustrating the translation of HCs, classified as memory or naive, into LCs predicted to belong to either maturation state. (C) Proportion of HCs for which *≥*80% of generated LCs utilize the same V gene family. Higher percentages indicate more constrained V gene family usage during LC generation. Match naive: heavy and LCs both predicted as naive; Match memory: heavy and LCs both predicted as memory; Mismatch naive: naive HC paired with memory-predicted LC; Mismatch memory: memory HC paired with naive-predicted LC.

To assess the structural quality of the generated sequences, the generated light sequences were folded using Chai-1 [41] and subsequently aligned with their reference sequences with ChimeraX [42] (Figure 2C). The generated LCs aligned well with their true paired counterparts, maintaining correct fold topology even at low sequence similarity (30.97%). These results suggest that our Heavy2Light model successfully captures essential structural constraints for antibody folding, prioritizing functiona fold preservation over exact sequence reproduction. The robust structural conservation across varied sequence similarities demonstrates the model’s ability to generate LCs that maintain the fundamental immunoglobulin framework.

To evaluate the biological compatibility of generated LCs with their corresponding HC inputs, we used ImmunoMatch [29], a protein language model that estimates the pairing probability between antibody chains. For each input HC, we generated 10 LC sequences and used our maturation state classifier to predict whether each sequence was memory or naive. Sequences were then grouped based on whether the predicted maturation state of the LC matched that of the input HC (Methods 4.5). Sequences with matching maturity states demonstrated superior pairing compatibility compared to non-matching pairs. When HCs and LCs shared the same predicted maturation state, the mean pairing score reached 0.6444, with 76.66% of sequences exceeding the threshold of 0.5. In contrast, sequences with non-matching maturity states achieved a markedly lower mean pairing score of 0.3708, with only 41.57% exceeding the compatibility threshold.

Among sequences with matched maturation states, naive-naive pairs achieved the highest average score (0.6490), followed closely by memory-memory pairs (0.6425). Despite these similar mean scores, memory-memory pairings displayed a distinct bimodal distribution, with a pronounced secondary peak around 0.9, indicating that a subset of memory-derived pairs achieved exceptionally high compatibility (Figure 3A). In contrast, mismatched pairs demonstrated lower compatibility. Surprisingly, naive HCs paired with memory LCs retained similar pairing ability to the matched maturation groups (average score: 0.6348), whereas memory HCs paired with naive LCs yielded the lowest compatibility (0.3118). While pairing scores clearly reflect the concordance between the predicted maturation states of the heavy and generated LCs, their correlation with sequence recovery, i.e., similarity to the reference LC, is weak for both groups (matching and mismatching maturity pairs, Supplementary Figure A7).

While naive HCs were predominantly paired with naive LCs, memory HCs showed a notable bias towards generating LCs predicted as naive. When the input HC was predicted as naive, on average 90.16% of the generated LC sequences were also predicted to be naive. In contrast, when HCs were predicted as memory B cells, only 48.10% of the generated LCs were predicted as memory (Figure 3B).

To find out more about the interdependence between our generated LCs and its input HCs, we evaluated the consistency of V gene family usage in generated LCs. For this, we stratified our data into four groups based on the predicted maturation states of HCs and LCs: (1) both predicted as naive, (2) both predicted as memory, (3) naive HCs paired with memory LCs, and (4) memory HCs paired with naive LCs (Methods section 4.6.2). Memory B cells exhibited greater V gene family constraint than naive B cells, with 16.1% of memory HCs showing focused V gene family usage (*≥*80% of generated LCs utilizing the same V gene family) compared to only 0.8% of naive HCs (Figure 3C). This pattern extended to mismatched pairings, where memory HCs paired with naive LCs maintained higher V gene constraint (11.7%) than naive HCs paired with memory LCs (5.3%). Similar V gene constraint patterns were observed across individual V genes as well (Supplementary Figure A8, Supplementary Table A6).

The conditioning strategy implemented in the Heavy2Light model not only strengthened sequence–structure relationships but also improved the biological plausibility of the generated LCs. When conditioned on a paired HC, the model produced LCs with high germline similarity across all antibody regions (Figure 4A, right panel). Framework regions retained high germline identity, with FR3 achieving 95.96%, FR1 at 95.01% and FR2 at 94.95%,. The CDRs achieved slightly lower germline similarity, with CDR2 at 85.76%, CDR3 at 85.62% and CDR1 at 78.88%. Overall, conditionally generated sequences achieved a total germline similarity of 93.10%. In comparison, unconditionally generated sequences (Figure 4A, left panel) showed slightly higher germline similarities, with framework regions consistently exceeding 95% (FR3 at 97.39%, FR2 at 96.39%, and FR1 at 95.68%) and CDRs ranging from 87.90% to 90.00%, resulting in a total germline similarity of 95.86%.

**Fig. 4.**
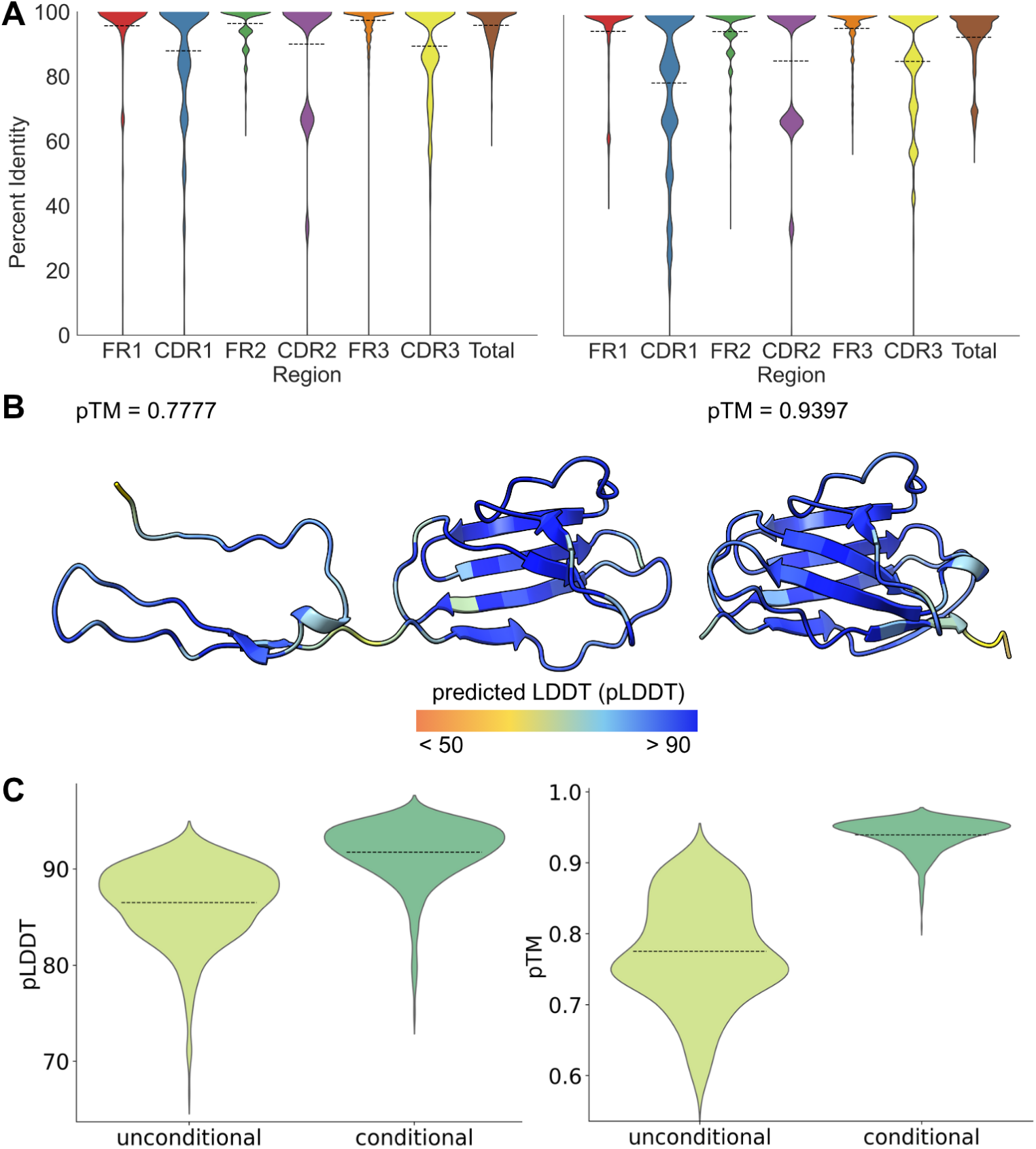
Comparison of unconditionally vs. conditionally generated LC sequences. (A) Germline identity across FR regions (FR1–FR3), CDRs (CDR1–CDR3), and complete variable domain. (B) Predicted structures for representive sequences. (For A and B: left panels show unconditional generation, right panels show conditional generation.) (C) Predicted structural quality metrics (pLDDT and pTM-score) for both generation modes.

Notably, the Heavy2Light model produced sequences with enhanced structural completeness compared to unconditioned generation. The total sequence length increased to a mean of 96.20 amino acids, representing a substantial improvement over the 80.59 amino acids observed in unconditioned sequences. This extension occurred across all regions: FR1 (25.52 vs. 20.84), FR2 (17.01 vs. 16.64), FR3 (36.00 vs. 32.21), CDR1 (7.39 vs. 6.87), CDR2 (3.03 vs. 3.02), and CDR3 (7.24 vs. 7.10) amino acids (Supplementary Table A7). The germline similarities as well as the mean lengths of each region and in total are similar to the distributions of sequence identity and mean lengths of the training and test set (Supplementary Figure A9, Supplementary Table A8). Importantly, structural folding revealed greatly improved folding quality for Heavy2Light-generated sequences (Figure 4B and C). The conditionally generated sequences achieved significantly higher predicted mean template modeling (pTM) scores (0.939) compared to unconditionally generated sequences (0.775), indicating substantially improved structural confidence. Similarly, predicted Local Distance Difference Test (pLDDT) scores were markedly higher for conditional generation (91.7) versus unconditional generation (86.5), suggesting more reliable local structural predictions throughout the sequence. Additionally, the generated LCs exhibited folding metrics comparable to their corresponding reference LCs, both when co-folded with their cognate heavy chains and when folded individually (Supplementary Figure A10 and Supplementary Table A9).

## 3 Discussion

Accurately modeling heavy–light chain pairing remains a central challenge in immunology, with broad implications for repertoire analysis, antibody discovery, and therapeutic engineering. Recent advances in deep learning have made it possible to extract meaningful patterns from large-scale immune sequence data. Here, we developed a two-stage deep learning framework: we first pre-trained two separate language models, HeavyBERTa for heavy chains and LightGPT for light chains, on large corpora of unpaired antibody sequences to learn chain-specific representations. These models were then integrated and fine-tuned on paired antibody data using lightweight adapters, enabling conditional generation of light chains from heavy chain input and providing a scalable approach to modeling inter-chain dependencies.

To investigate functional constraints on antibody pairing, we trained classifiers to distinguish between naive and memory B cell sequences based on primary sequence alone. The heavy chain classifier consistently outperformed its light chain counterpart, likely due to the greater sequence diversity in HCs introduced by VDJ recombination and junctional variability [2, 4–6]. To evaluate the biological plausibility of these predictions, we applied the classifier to unlabeled paired datasets and assessed V gene coherence across the predicted maturation states. Sequences classified as memory showed greater V gene coherence between HC and LC than those classified as naive, echoing patterns previously reported in experimentally annotated datasets [34]. Interestingly, the frequency of coherent pairings was even higher than previously reported, probably reflecting dataset biases, such as restricted donor representation and the under-sampling of naive compartments [14, 35]. Memory-classified heavy chains generated light chains with restricted V gene usage, even when the generated sequences were classified as naive. In contrast, naive-classified HCs yielded LCs with broader V gene diversity, consistent with a more permissive, less selection-constrained repertoire. These results suggest that maturation imposes directional constraints on pairing preferences, which can be learned and reproduced by a generative model.

Despite the modest sequence recovery of true light chains by our Heavy2Light model, we observed that many generated sequences still achieved high Immuno-Match scores—suggesting that functional compatibility may be preserved even when exact sequence matches are not. This weak correlation between recovery rate and ImmunoMatch compatibility implies that the model captures key features important for pairing beyond primary sequence identity. Prior studies have shown that specific residues at defined positions can determine heavy–light chain pairing modes [43], and that point mutations in CDR-H3 or CDR-L3 can reshape VH–VL interfaces through conformational changes without major sequence shifts [44]. These findings suggest that generated light chains may retain essential structural or functional determinants for pairing, even when they diverge from the native sequence.

The trimodal distribution of sequence similarity observed in generated *κ* light chains across both naive and memory groups could not be explained by V gene usage bias. We propose that this pattern reflects a biological organization within the *κ* light chain repertoire, presenting distinct challenges for sequence prediction. The lowest similarity mode may represent highly promiscuous, largely public light chains capable of pairing with diverse heavy chains [16], while the higher similarity modes likely correspond to more specialized or antigen-driven variants that have co-evolved with their cognate heavy chains. This spectrum suggests a continuum from broadly compatible, promiscuous light chains to increasingly specific, co-adapted HC-LC pairings. In contrast, *λ* light chains exhibited a dominant low-recovery mode, consistent with previous reports that *λ* chains have greater structural flexibility, primarily due to a longer and more diverse CDR-L3 region [45] and increased amino acid diversity in both framework and CDR regions compared to *κ* chains [46]. This expanded conformational and sequence variability likely complicates accurate sequence prediction. Moreover, the VL domain in *λ* chains is less stabilized, and VL–VH interactions differ from those in *κ* chains [45], potentially leading to less stereotyped pairing patterns and further hindering inference of light chain sequences solely from heavy chain context.

The complex and heterogeneous patterns of sequence similarity in generated light chains underscore the challenges of accurately modeling antibody pairing. Despite this, conditioning light chain generation on heavy chain input markedly improved the structural and biological plausibility of the outputs. Conditioned sequences exhibited more complete antibody regions, with improved coverage of both framework and CDR segments. Structure-based metrics further supported these gains, with conditioned light chains achieving higher pLDDT and pTM scores compared to those generated without heavy chain context. These improvements suggest that the Heavy2Light model captures inter-chain dependencies relevant to productive antibody folding, even when sequence-level recovery is modest. While experimental validation remains necessary to confirm functional relevance, these results point to the potential of conditional generative modeling as a tool for exploring and designing biologically coherent antibody pairs.

Our work support an emerging paradigm in which antibody sequence modeling moves beyond classification or annotation toward generative biology, where models propose novel, yet biologically grounded, sequences. We observed that coherence in predicted maturation states between heavy and light chains was associated with more compatible pairings, suggesting that maturation-aware generation contributes to biologically plausible outputs. However, the model exhibited a tendency to produce naive-like light chains, reflecting known limitations of current antibody datasets and language models trained on germline-skewed repertoires [47]. This bias likely arises from the prevalence of germline-proximal sequences in public datasets, driven both by the dominance of naive B cells in peripheral blood and by the fact that even affinity-matured antibodies often retain high germline identity. These imbalances pose challenges for training models that aim to capture the full diversity of affinity-matured repertoires. Addressing this will require more balanced datasets, including memory and therapeutic sequences, along with advances at the model level, such as incorporating structural priors, contrastive objectives, or alternative generative frameworks such as diffusion models. Looking forward, extending generative modeling beyond structural compatibility toward functional endpoints—such as antigen specificity, epitope targeting, or neutralization—will be essential for translational applications. Such models also hold promise for interrogating repertoire dynamics in infection, vaccination, or autoimmunity, opening new avenues for systems-level immunological insights.

This work demonstrates the feasibility of using deep learning to model the interdependent sequence features that govern antibody pairing and maturation. By integrating discriminative and generative approaches, we show that transformer-based architectures can uncover immunologically meaningful constraints and enable conditional antibody design. These findings lay the foundation for future tools that support repertoire analysis, therapeutic engineering, and the broader goals of data-driven immunology.

## 4 Methods

### 4.1 Models

We pre-trained two antibody-specific language models based on the transformer [48] architecture: HeavyBERTa for HC sequences and LightGPT for LC sequences. For HeavyBERTa, we implemented two RoBERTa-based configurations to evaluate the impact of model size on performance. The small configuration featured a hidden size of 512, an intermediate size of 2048, 4 attention heads, and 4 transformer layers, totaling approximately 13.15 million parameters. We then scaled up to a larger configuration with a hidden size of 768, an intermediate size of 3072, 12 attention heads, and 12 transformer layers, resulting in approximately 86.06 million parameters. For LightGPT, we used a GPT-2 architecture with a hidden size of 768, 12 attention heads, and 12 transformer layers, totaling approximately 85.86 million parameters. Both architectures utilized a vocabulary of 25 tokens, comprising 20 standard amino acids plus 5 special tokens. For downstream applications, we employed parameter-efficient fine-tuning using bottleneck adapters [40] with multi-head and output adapter configurations enabled. The adapters utilized a reduction factor of 16 and Rectified Linear Unit (ReLU) as activation function (Supplementary Table A10). The Heavy2Light encoder-decoder model combined the pre-trained HeavyBERTa (encoder) and LightGPT (decoder) architectures with cross-attention mechanisms, where only adapter modules and cross-attention parameters remained trainable while pre-trained weights were frozen (Figure 1). Similarly, for the classification of HC and LC sequences into naive or memory B cell states, we fine-tuned the respective pre-trained models (Heavy-BERTa for HCs, LightGPT for LCs) using the same adapter configuration, adding classification heads while keeping the base model parameters frozen.

### 4.2 Data preparation

All models were trained using data from the Observed Antibody Space (OAS [35]). Additionally, paired data from the Patent and Literature Antibody Database (PLAbDab [39]) was used for the Heavy2Light fine-tuning dataset. Four different datasets were created: a HC dataset for HeavyBERTa pre-training, a LC dataset for LightGPT pre-training, a paired sequences dataset for Heavy2Light fine-tuning and another paired dataset for the B cell classification task.

#### 4.2.1 Pre-training datasets for HeavyBERTa and LightGPT

For the HeavyBERTa and LightGPT pre-training datasets, unsorted B cells from peripheral blood mononuclear cells (PBMCs), memory B cells, and plasma B cells were selected. For the OAS data, we selected only sequences from healthy donors without disease or vaccination history. Additionally, we incorporated all available paired sequences from the PLAbDab database. To reduce sequence redundancy and ensure proper data splits, we implemented a hierarchical clustering strategy for the pre-training datasets. First, CDR3 sequences (CDRH3 for HCs, CDRL3 for LCs) from the OAS database were clustered at 100% sequence identity using MMseqs2’s linclust module (version 13.45111) [49]. The centroids from these clusters were then used to extract full-length HC or LC sequences, which were subsequently clustered at 70% sequence identity. These 70% identity clusters formed the basis of our training datasets. To prevent highly similar sequences from appearing in both training and test sets, we performed an additional clustering step at 50% sequence identity and allocated sequences to training, validation, and test datasets based on clusters rather than random assignment. This approach ensures that sequences within the same 50% identity cluster remain together in the same dataset split, thereby preventing data leakage and providing more robust model evaluation.

#### 4.2.2 Fine-tuning dataset for Heavy2Light

For the Heavy2Light fine-tuning dataset, we combined paired sequences from both OAS (human, healthy donors without vaccination or disease) and PLAbDab databases. After duplicate removal, the paired sequences were clustered at 30% sequence identity for allocation into training (80%), testing (10%), and validation (10%) splits, following the same cluster-based allocation strategy to maintain sequence independence across datasets.

#### 4.2.3 Fine-tuning dataset for naive/memory B cell classification

To construct a balanced and non-redundant dataset of naive and memory B cell receptor sequences, we started with all the paired data sequences available from OAS. For each antibody chain (heavy and light) separately, we extracted the aligned amino acid sequences and filtered them to retain unique entries. To balance the dataset, we randomly sampled an equal number of Naive and Memory B cell sequences, using the smaller of the two class sizes as reference. To minimize sequence redundancy across data splits, we clustered the sequences using linclust with default parameters, processing HCs and LCs separately. Clustering information was then used to perform non-overlapping train, validation, and test splits: all sequences from the same cluster were assigned to the same subset. This strategy mitigates information leakage across splits. The final dataset was divided into training (80%), validation (10%), and test (10%) sets, with larger clusters preferentially assigned to the training set.

### 4.3 Pre-training and fine-tuning

#### 4.3.1 HeavyBERTa pre-training

Two RoBERTa-based configurations were pre-trained on unpaired HC sequences using masked language modeling. Both models were trained with identical hyperparameters: a learning rate of 5e-5, weight decay of 0.1, batch size of 16, and the AdamW optimizer with linear learning rate scheduling. The small configuration (4 layers, 512 hidden size) was trained for 22 epochs over 136,254,500 training steps, requiring approximately 244 hours of computation. The large configuration (12 layers, 768 hidden size) was trained for 9.18 epochs over 56,832,000 training steps, requiring approximately 673 hours (for more information, see Supplementary Table A11).

#### 4.3.2 LightGPT pre-training

The GPT-2-based model was pre-trained on unpaired LC sequences using autoregressive language modeling. Training was conducted for 41 epochs over 12,265,000 steps with a batch size of 16, learning rate of 5e-5, weight decay of 0.1, and AdamW optimization with linear scheduling. The model required approximately 606 hours of training time (Supplementary Table A12).

#### 4.3.3 Fine-tuning Heavy2Light encoder-decoder model

The Heavy2Light translation model was fine-tuned using parameter-efficient adapter-based training, combining the pre-trained HeavyBERTa encoder (small configuration) with the LightGPT decoder in an encoder-decoder architecture. Training was conducted for 50 epochs with a batch size of 64, learning rate of 1e-5, weight decay of 0.1, and gradient clipping with a maximum norm of 1.0. We used AdamW as optimizer with linear learning rate scheduling. For sequence generation, nucleus sampling was implemented with a top-p value of 0.85, temperature of 0.8, and top-k disabled (set to 0). Generation length was constrained with a maximum of 115 new tokens. For each HC, 10 LCs were generated. For the germline similarities and the true LC recoery, we extracted the first generated light sequence which matched the maturity of the given input HC. For the other analyses, we used all generated LCs. The model was trained for 367,750 steps over approximately 34.9 hours (Supplementary Table A10).

#### 4.3.4 Classification model fine-tuning

Classification models for naive/memory B cell prediction were fine-tuned using the same adapter-based approach on their respective pre-trained base models. The Heavy-BERTa classifier was trained for 200 epochs with a learning rate of 3e-6, batch size of 64, maximum sequence length of 150, and dropout rate of 0.1. The LightGPT classifier was trained for 50 epochs with an identical learning rate and batch size but with a higher dropout rate of 0.3. Both models utilized AdamW optimization with weight decay of 0.01 and maintained frozen base model parameters while training only the adapter modules and classification heads (Supplementary Table A13).

### 4.4 Evaluation

The generated light sequences were evaluated using alignment-based percentage similarity. The alignment of the generated and true sequences was done with global alignment using Biopython (version 1.79), with a gap opening penalty of −10 and a gap extension penalty of −4. Chai-1 (version 0.6.1) [41] was used for protein structure predictions of the generated and true LCs using multiple sequence alignments. The generated and corresponding true LC were aligned, superimposed and visualised using ChimeraX (version 1.9) [42]. All dimensionality reduction visualizations were generated with the following parameters and configurations: Embeddings were extracted from the final hidden layer of each pre-trained model using mean pooling across sequence positions. PCA, t-SNE, and LDA were performed using scikit-learn (version 1.5.1). T-distributed Stochastic Neighbor Embedding (t-SNE) was applied with a perplexity of 30. LDA supports a maximum of *n −* 1 components for *n* classes, so we included three groups (memory, naive, RV+B cells) for dimensionality reduction (Supplementary Figure A4), but visualized only the naive and memory groups. PyIR [50], a wrapper for the IgBLAST [51] immunoglobulin and T-cell analyser, was used to calculate the percent identity to the germline.

### 4.5 Pairing compatibility assessment using ImmunoMatch

To assess the biological compatibility of generated LCs with their corresponding HC inputs, we applied the recently published ImmunoMatch model [29], which estimates pairing probabilities based on sequence features. For each input HC, we generated 10 LCs and predicted their maturation state (naive or memory) using our classification model. Pairings were initially grouped based on maturation state concordance between the HC and generated LCs: (1) Matching (both predicted as naive or both as memory), and (2) Non-matching (discordant maturation predictions). For further analysis, these were further stratified into four subgroups: (1) Matching memory (both HC and LCs predicted as memory), (2) Matching naive, (3) Non-matching memory (memory HC with naive LCs), and (4) Non-matching naive (naive HC with memory LCs). Prior to ImmunoMatch evaluation, all sequences were standardized using PyIR [50] to trim the variable regions and remove any non-canonical residues.

### 4.6 HC-LC V gene interdependence

#### 4.6.1 V gene consistency in OAS database

To investigate HC-LC interdependence patterns, we analyzed paired sequences from the OAS database. We grouped HCs based on their CDRH3 regions and their V gene. We then removed all groups containing only single entries and excluded groups consisting of sequences from only one patient [34]. For coherence calculation, we applied our trained HeavyBERTa classifier to predict B cell origin (naive or memory) for HC sequences in paired datasets. For each group containing more than one sequence and originating from different patients, we determined whether the LC V gene was identical across all sequences sharing the same HC V gene. Coherence was calculated as the fraction of groups showing identical LC V gene usage within each predicted B cell category.

#### 4.6.2 V gene consistency of conditionally generated light sequences

To assess V gene family usage consistency in generated LCs, we performed a frequency-based analysis across four maturation state pairings between heavy and LCs: (1) both predicted as naive, (2) both predicted as memory, (3) naive HC with memory LCs, and (4) memory HC with naive LCs. For each HC in the dataset, ten LC sequences were generated using our model. Each generated LC was annotated with its V gene and V gene family using PyIR [50]. We then grouped the generated LCs by their corresponding input HC (i.e., all LCs generated from the same HC sequence) and calculated the frequency distribution of V genes and V gene families within each group. For each HC group, we recorded the proportion of sequences sharing the most frequently used V gene or family. A HC was considered to exhibit V gene constraint if *≥*80% of its associated LCs used the same V gene or V gene family. This approach allowed us to quantify the extent of V gene restriction across different maturation-state pairings. The *≥*80% threshold was chosen to identify cases where V gene usage was strongly focused. Only HCs with a minimum of four associated generated LCs were included.

### 4.7 Hardware and software

All models were trained using the HuggingFace Transformers library (version 4.40.2) [48]. We used the Adapters (version 1.0.0.dev0) [40] library for the encoder-decoder models to facilitate modular fine-tuning. PyTorch was used as backend (version 2.3.1) [52] for both the HuggingFace Transformers Library and the Adapters library. Pre-training and fine-tuning were conducted on NVIDIA A100 or H100 Tensor Core GPUs (each with 80GB of RAM), depending on availability.

## 5 Resource availability

All code, pretrained models, and curated datasets used in this study are openly available to support transparency, reproducibility, and community-driven development. Resources can be accessed via GitHub at the following repository: https://github.com/ibmm-unibe-ch/Heavy2Light

## 6 Acknowledgments

The project was supported by the Helmut Horten Young Investigator Program 2022 (project ID: 2022-YIG-089).

## 7 Author contributions

LB acquired and analyzed data, designed and trained models, and drafted the manuscript with critical revision and final approval. TL contributed to study design, article drafting, critical revision, and final approval. CR conceived the original idea and participated in study design, data analysis, article drafting, critical revision, and final approval of the manuscript.

## 8 Declaration of interests

The authors declare no competing interests.

### 8.1 Declaration of generative AI and AI-assisted technologies in the writing process

During the preparation of this manuscript, the authors employed generative AI tools, including ChatGPT (OpenAI) and Gemini (Google), for language refinement and editorial assistance. These tools were used solely to improve clarity, coherence, and style. All AI-assisted content was carefully reviewed and substantively edited by the authors, who accept full responsibility for the accuracy and integrity of the final submitted work.

## Appendix A Supporting Information

**Fig. A1.**
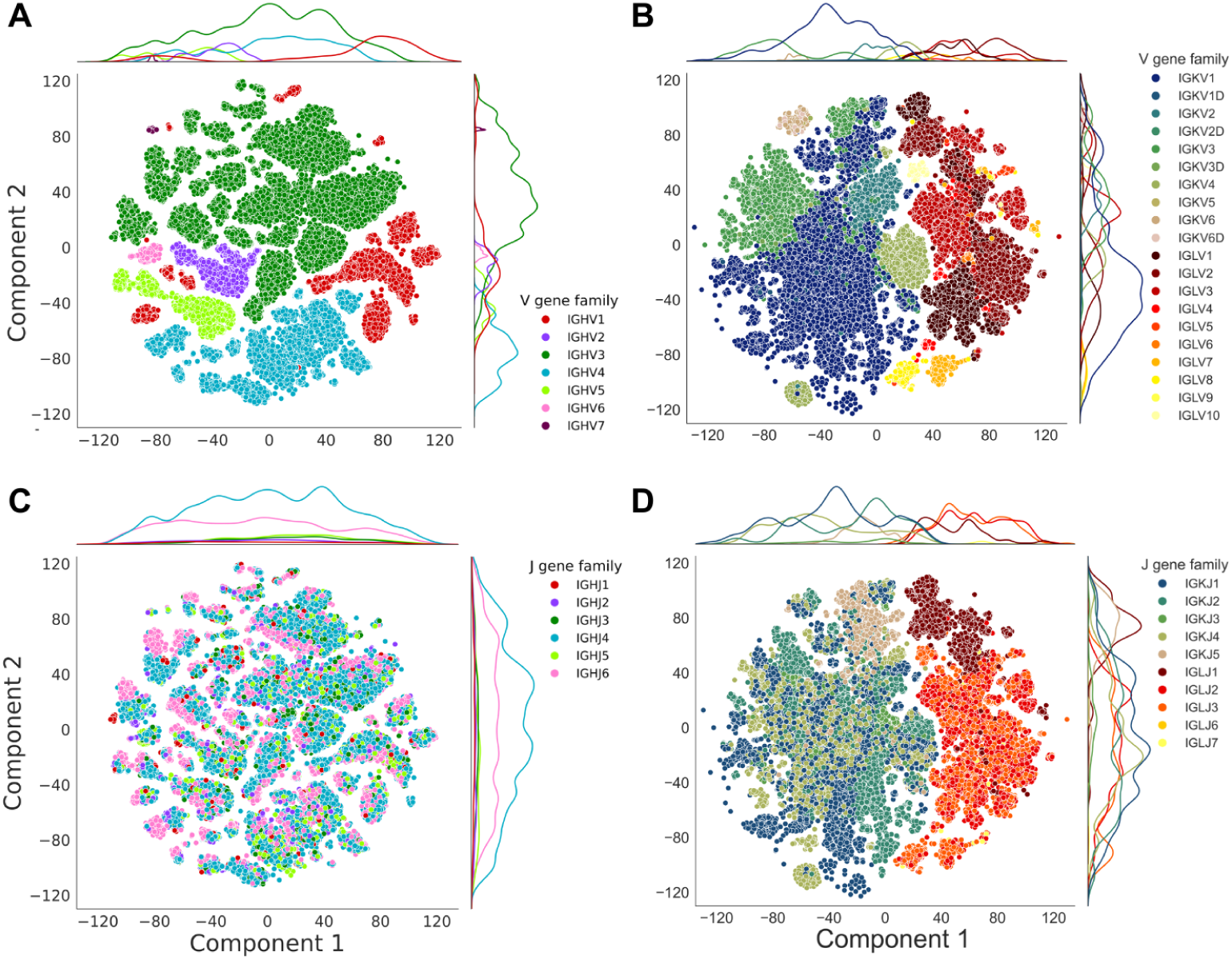
T-SNE visualization of final layer embeddings of V (A, B) and J (C, D) gene families for both HeavyBERTa (A, C) and LightGPT (B, D) models. Each point represents an individual sequence, colored by V or J gene family assignment (*κ* gene families displayed in cool colors and *λ* gene families in warm colors). Density plots along each axis show the distribution of sequences across principal components.

**Fig. A2.**
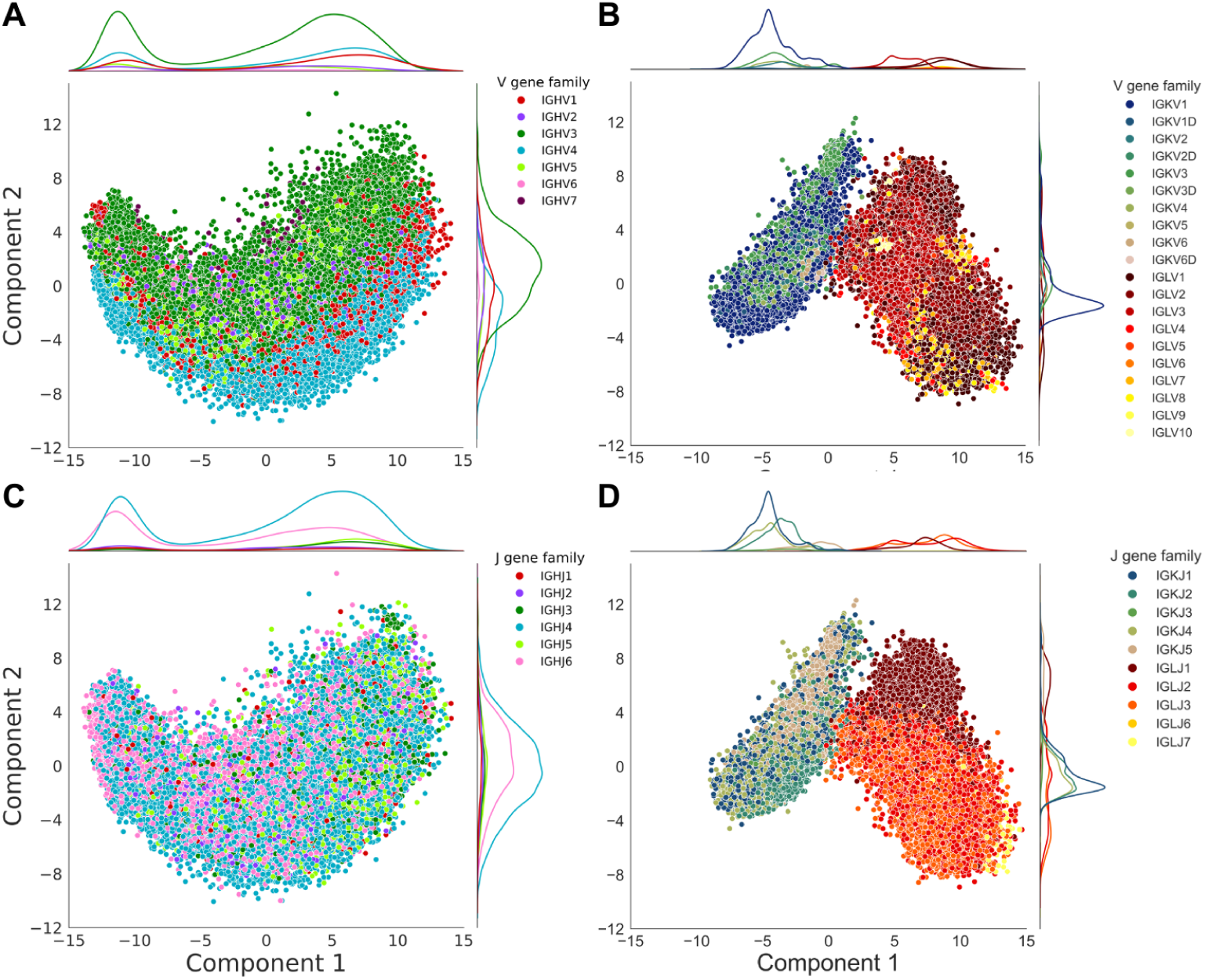
PCA of learned embeddings colored by gene family annotations. HeavyBERTa (A, C) embeddings projected onto the first two principal components, with sequences colored by HC V gene families (A, B) and J gene families (C, D). LightGPT (B, D) embeddings projected onto the first two principal components, with sequences colored by LC V gene families (top) and J gene families (bottom). Each point represents an individual sequence (*κ* gene families displayed in cool colors and *λ* gene families in warm colors). Density plots along each axis show the distribution of sequences across principal components.

**Fig. A3.**
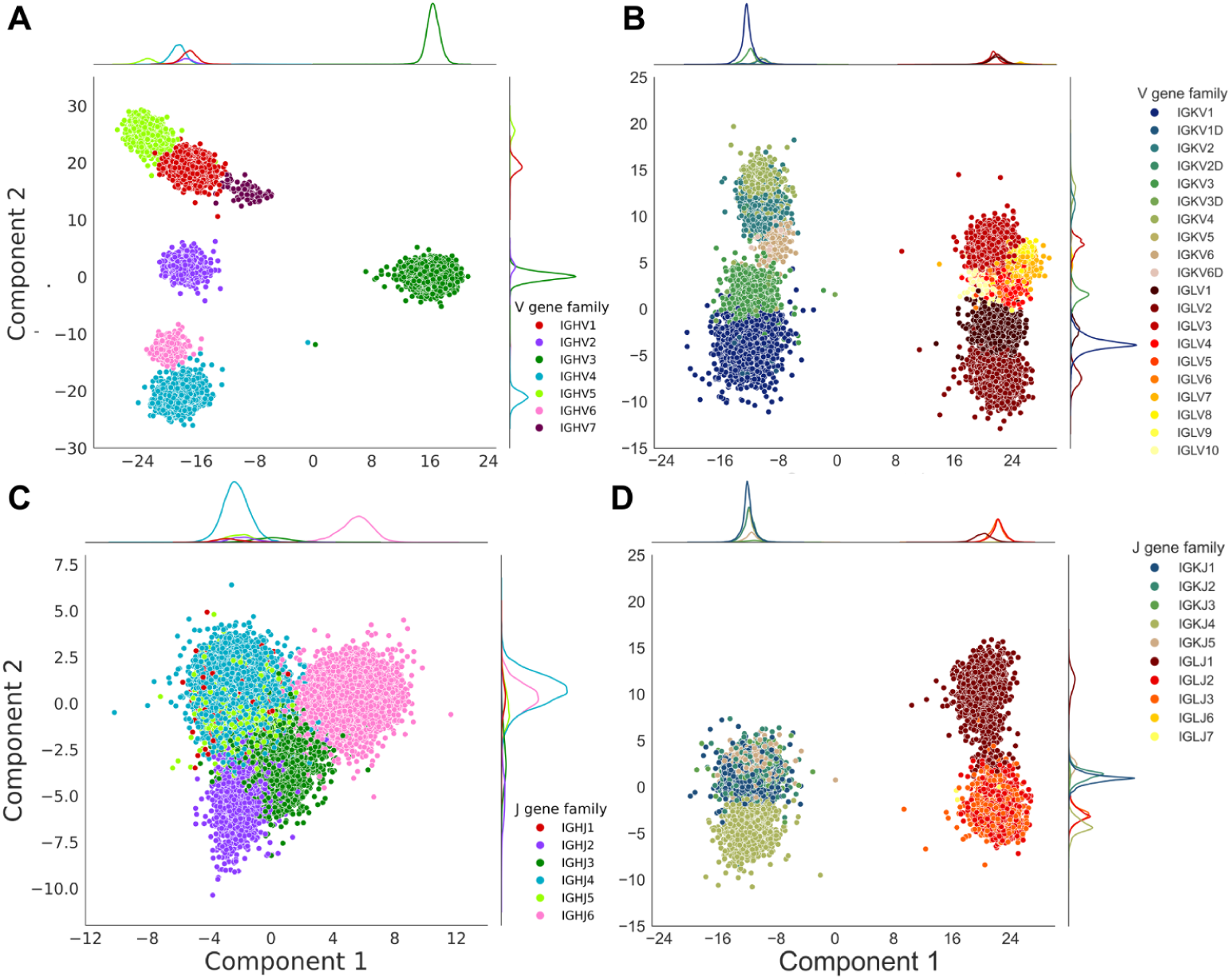
LDA of learned embeddings colored by gene family annotations. HeavyBERTa (A, C) embeddings projected onto the first two linear discriminants, with sequences colored by HC V gene families (A) and J gene families (C). LightGPT (B, D) embeddings projected onto the first two linear discriminants, with sequences colored by LC V gene families (B) and J gene families (D). Each point represents an individual sequence (*κ* gene families displayed in cool colors and *λ* gene families in warm colors). Density plots along each axis show the distribution of sequences across linear discriminants.

**Fig. A4.**
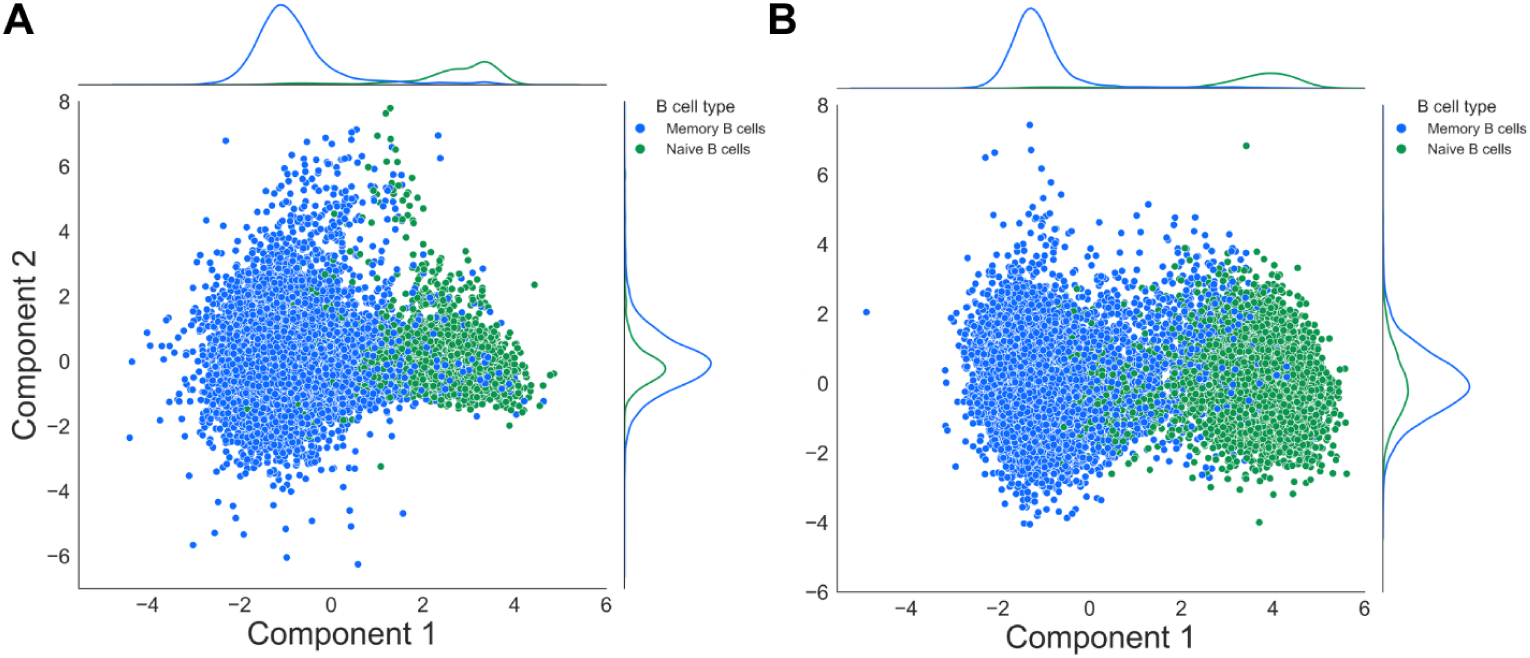
LDA of final layer embeddings. LDA effectively separates memory and naive B cell populations for HeavyBERTa (B), as well as LightGPT (D), indicating that the learned representations encode biologically relevant features associated with B cell developmental states.

**Table A1.**
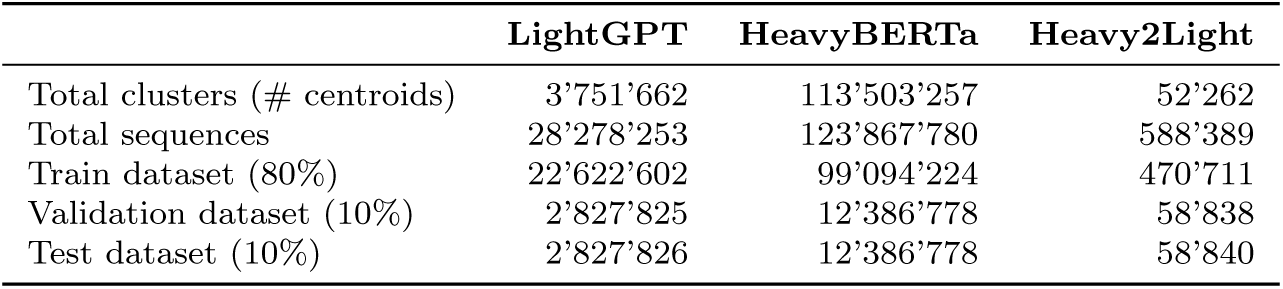
Dataset composition and data split allocation. . Number of sequences and centroids used for training, validation, and testing across all model datasets. HeavyBERTa and LightGPT datasets show unpaired sequence counts with corresponding centroids used for cluster-based allocation (50% identity clustering). The Heavy2Light dataset shows paired sequence counts from combined OAS and PLAbDab sources with centroids derived from 30% identity clustering for allocation across data splits.

|  | LightGPT | HeavyBERTa | Heavy2Light |
| --- | --- | --- | --- |
| Total clusters (# centroids) | 3'751'662 | 113'503'257 | 52'262 |
| Total sequences | 28'278'253 | 123'867'780 | 588'389 |
| Train dataset (80%) | 22'622'602 | 99'094'224 | 470'711 |
| Validation dataset (10%) | 2'827'825 | 12'386'778 | 58'838 |
| Test dataset (10%) | 2'827'826 | 12'386'778 | 58'840 |

**Table A2.** Language modeling performance across model configurations. . Accuracy values represent the fraction of correctly predicted amino acid residues for masked language modeling (HeavyBERTa models with 15% random masking) and autoregressive language modeling (LightGPT model with next-token prediction), evaluated on their respective HC and LC test datasets.

| Model | Configuration | Antibody chain type | Accuracy |
| --- | --- | --- | --- |
| RoBERTa | small | Heavy | 0.8895 |
| RoBERTa | large | Heavy | 0.8912 |
| GPT-2 | – | Light | 0.8635 |

**Table A3.**
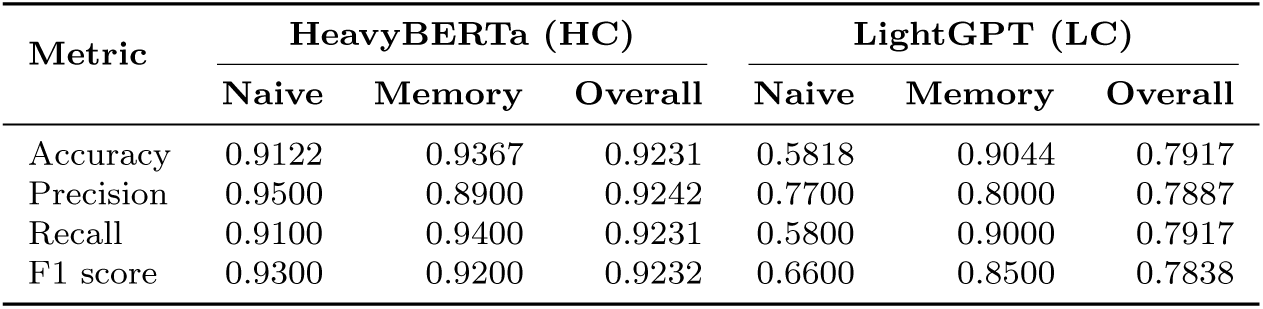
Classification performance for naive and memory B cell prediction. . Performance metrics for HeavyBERTa and LightGPT models in classifying antibody sequences by B cell developmental state. Metrics are reported for individual classes (naive and memory) as well as overall performance across the entire test dataset.

**Table A4.** Coherence percentages of V gene usage. Coherence percentages of V gene usage in memory and naive B cells for the labeled OAS database data and the previously unlabeled data classified by our model.

| Dataset | Memory B cells | Naive B cells |
| --- | --- | --- |
| OAS | 0.55817 | 0.13834 |
| Classified from unsorted B cells | 0.80571 | 0.05595 |
| All combined data | 0.60634 | 0.11190 |

**Table A5.**
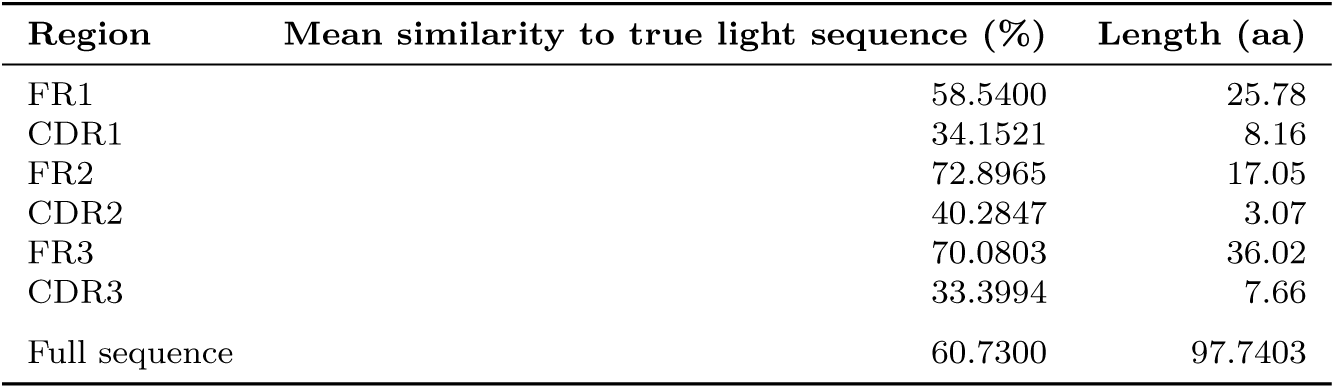
Similarity of generated LC sequences to their corresponding true light sequence. . Comparison of identity percentages across antibody regions (FR1–3, CDR1–3, and total sequence) for LCs generated conditionally by Heavy2Light. Values represent mean germline similarity scores to its true light sequence after global alignment; Length indicates the average region length in amino acids.

**Table A6.** Numbers and sample sizes for V gene constraint analysis showing proportion of HCs with *≥*80% V gene consistency in generated LCs. Match naive: heavy and LCs both predicted as naive; Match memory: heavy and LCs both predicted as memory; Mismatch naive: naive HC paired with memory-predicted LC; Mismatch memory: memory HC paired with naive-predicted LC.

|  | Match Naive |  | Match Memory |  | Mismatch Naive |  | Mismatch Memory |  |
| --- | --- | --- | --- | --- | --- | --- | --- | --- |
|  | Count | % | Count | % | Count | % | Count | % |
| Full V gene | 8/13,864 | 0.1 | 1,478/25,694 | 5.8 | 16/392 | 4.1 | 338/29,321 | 1.2 |
| V gene family | 114/13,864 | 0.8 | 4,135/25,694 | 16.1 | 46/392 | 11.7 | 1,548/29,321 | 5.3 |

**Table A7.** Germline similarity and recognized sequence length of generated LC sequences. Comparison of germline identity percentages and amino acid lengths across antibody regions (FR1-3, CDR1-3, and total sequence) for LCs generated unconditionally by LightGPT versus conditionally by the Heavy2Light model. Values represent mean germline similarity scores and sequence lengths in amino acids as determined by IgBLAST alignment analysis.

| Region | Germline similarity |  | Length [AA] |  |
| --- | --- | --- | --- | --- |
|  | LightGPT | Heavy2Light | LightGPT | Heavy2Light |
| FR1 | 0.9568 | 0.9501 | 20.84 | 25.52 |
| CDR1 | 0.8790 | 0.7888 | 6.87 | 7.39 |
| FR2 | 0.9639 | 0.9495 | 16.64 | 17.01 |
| CDR2 | 0.9000 | 0.8576 | 3.02 | 3.03 |
| FR3 | 0.9739 | 0.9596 | 32.21 | 36.00 |
| CDR3 | 0.8934 | 0.8562 | 7.10 | 7.24 |
| Full sequence | 0.9586 | 0.9310 | 80.59 | 96.20 |

**Table A8.** Germline identity across antibody regions (FR1–FR3, CDR1–CDR3, and full sequence) for LCs in the training, validation, and test sets. Sequence identity is expressed as the fraction of germline-matched residues; Length is the average region length in amino acids.

| Region | Training |  | Validation |  | Test |  |
| --- | --- | --- | --- | --- | --- | --- |
|  | Seq. Id. | Length | Seq. Id. | Length | Seq. Id. | Length |
| FR1 | 0.9565 | 25.59 | 0.9310 | 25.39 | 0.9348 | 25.39 |
| CDR1 | 0.8595 | 7.48 | 0.7536 | 7.84 | 0.7510 | 7.22 |
| FR2 | 0.9593 | 17.01 | 0.9099 | 17.03 | 0.9123 | 17.02 |
| CDR2 | 0.8944 | 3.05 | 0.7749 | 3.06 | 0.7958 | 3.03 |
| FR3 | 0.9652 | 36.03 | 0.9279 | 36.03 | 0.9278 | 36.01 |
| CDR3 | 0.8902 | 7.26 | 0.7960 | 7.33 | 0.8049 | 7.04 |
| Full sequence | 0.9448 | 96.41 | 0.8967 | 96.66 | 0.9000 | 95.64 |

**Fig. A5.**
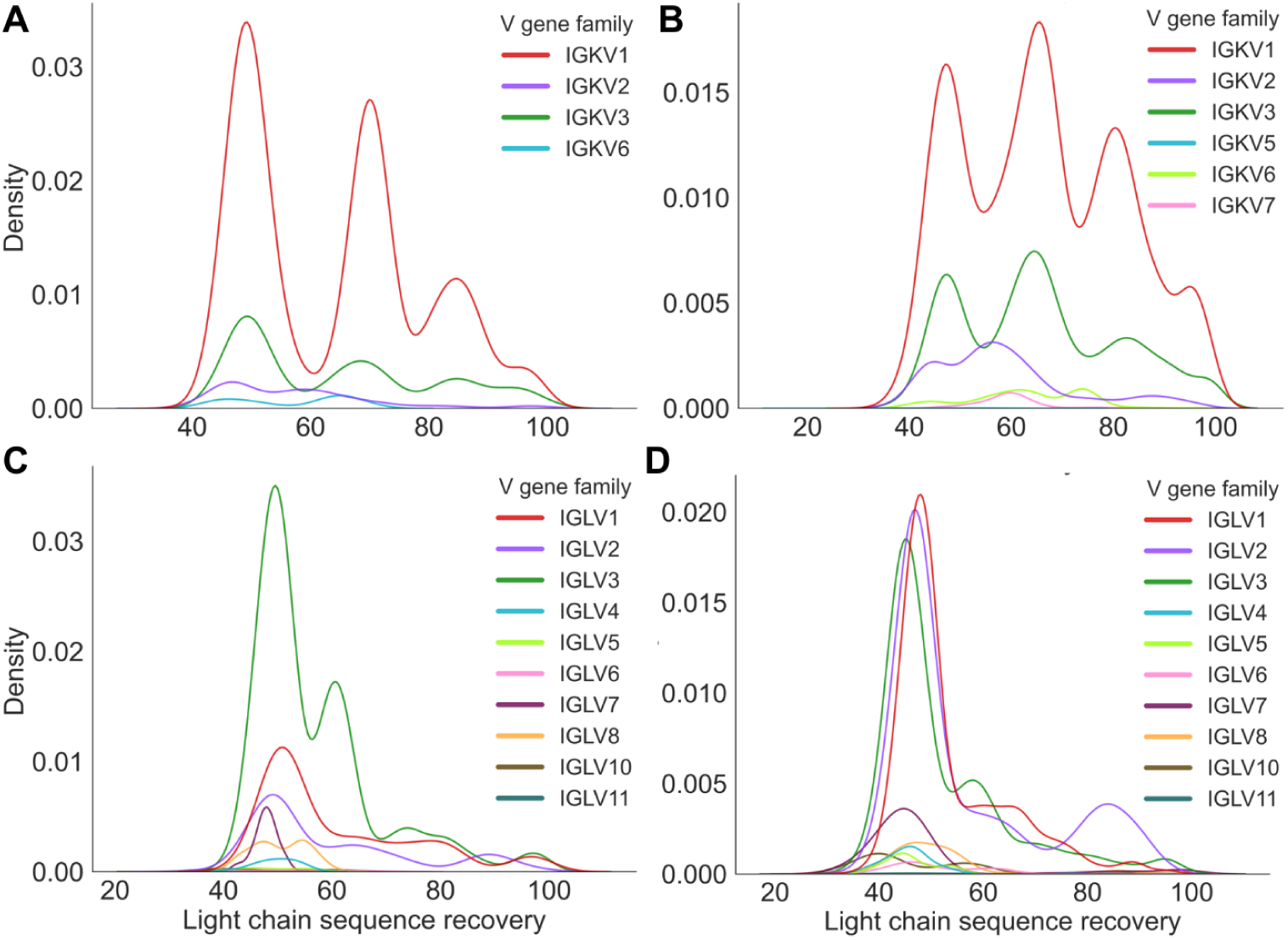
LC sequence recovery distributions by V gene family across B cell subsets and LC types. Kernel density plots showing the distribution of percent identity between predicted and true LC sequences, grouped by V gene family. (A) Kappa LCs from naive B cells, (B) Kappa LCs from memory B cells, (C) Lambda LCs from naive B cells, and (D) Lambda LCs from memory B cells.

**Fig. A6.**
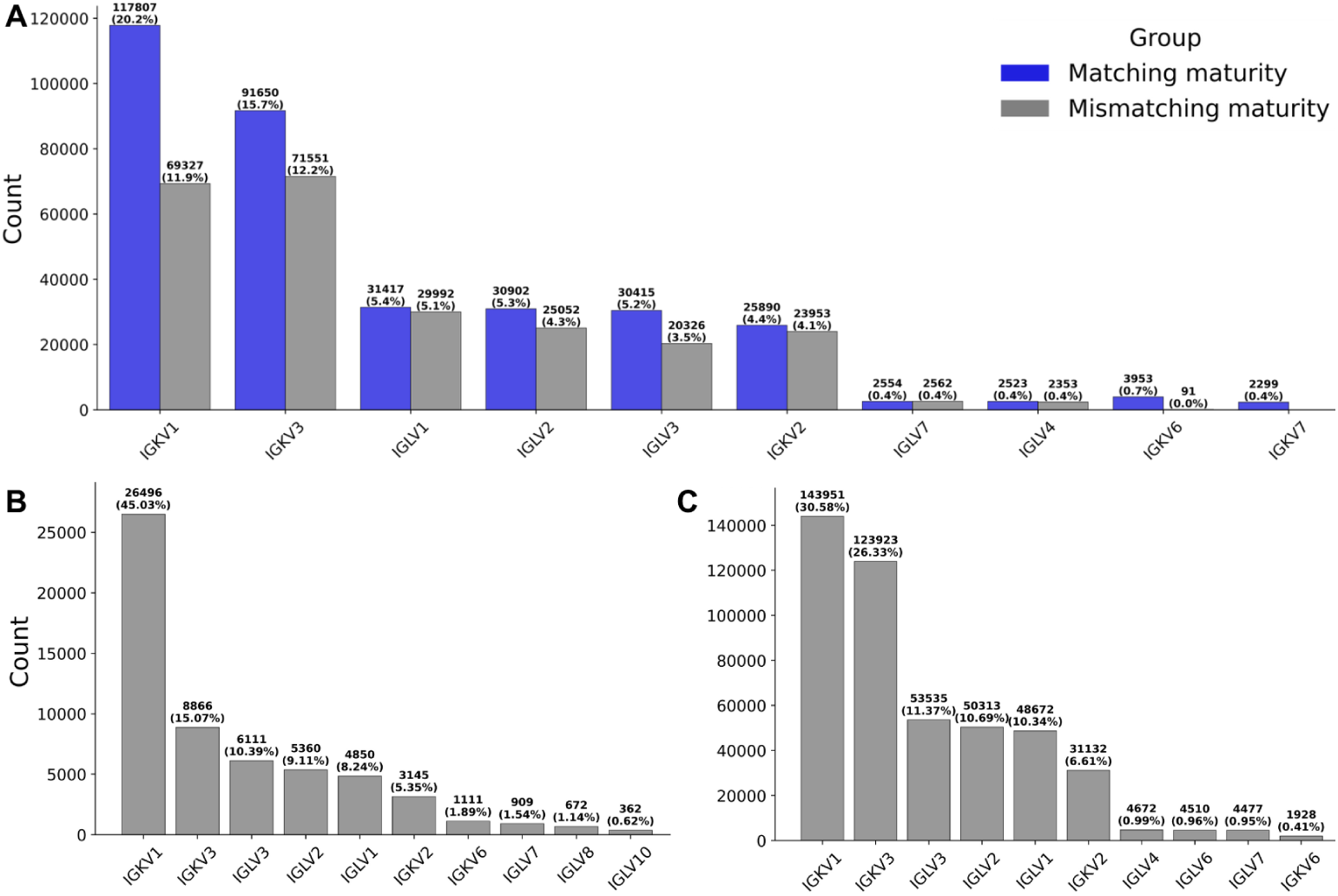
V gene family distribution across generated and reference antibody LC sequences. (A) V gene family usage in all generated sequences from the Heavy2Light model, (B) true LC sequences from the test set, and (C) true LC sequences from the training set. IGKV1 predominates across all datasets. Numbers above bars indicate absolute counts and percentages.

**Fig. A7.**
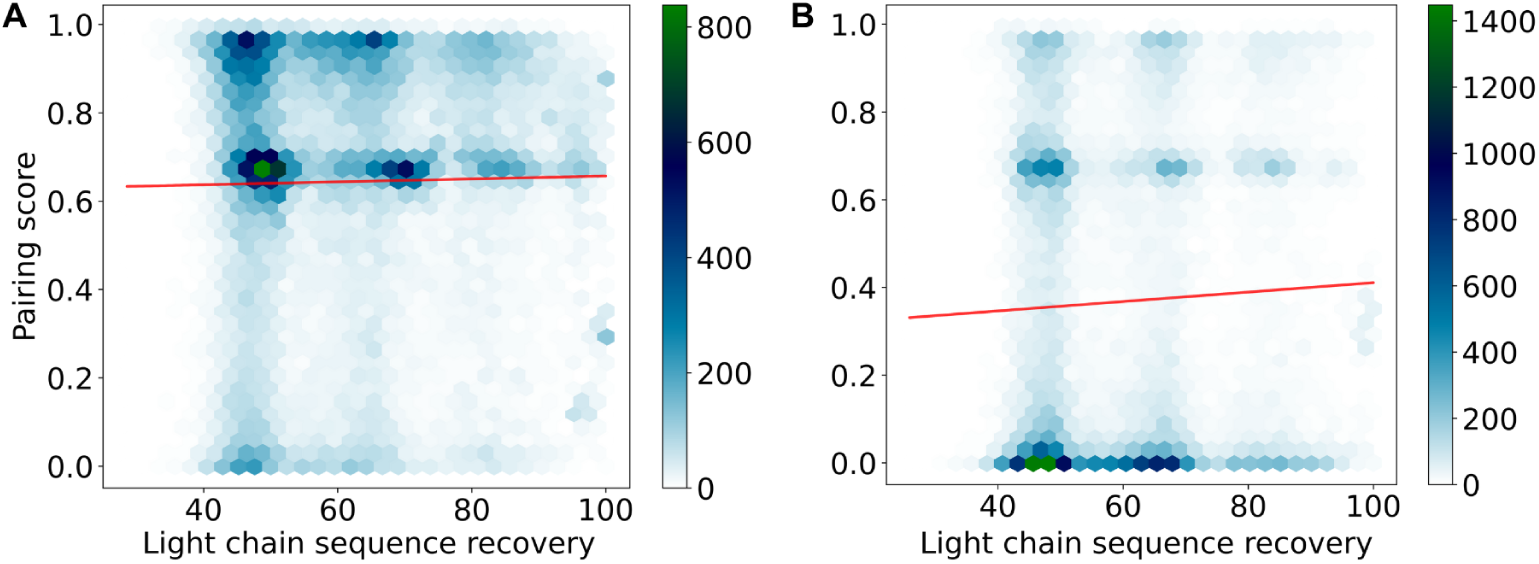
Correlation between ImmunoMatch [29] pairing probability scores and sequence recovery rates. Scatter plots showing the relationship between ImmunoMatch pairing probability scores and sequence identity for (A) maturity-matched heavy-LC pairs and (B) maturity-mismatched HC-LC chain pairs. Matched pairs correspond to heavy and generated LCs both predicted to have the same maturation state (memory or naive), while mismatched pairs have different predicted maturation states between heavy and generated LCs. Maturity-matched pairs show a weak positive correlation (Pearson correlation coefficient *r* = 0.0184, *p* = 1.290 *×* 10*^−^*^5^), while maturity-mismatched pairs exhibit a slightly stronger but still weak correlation (*r* = 0.0461, *p* = 9.653*×*10*^−^*^24^).

**Fig. A8.**
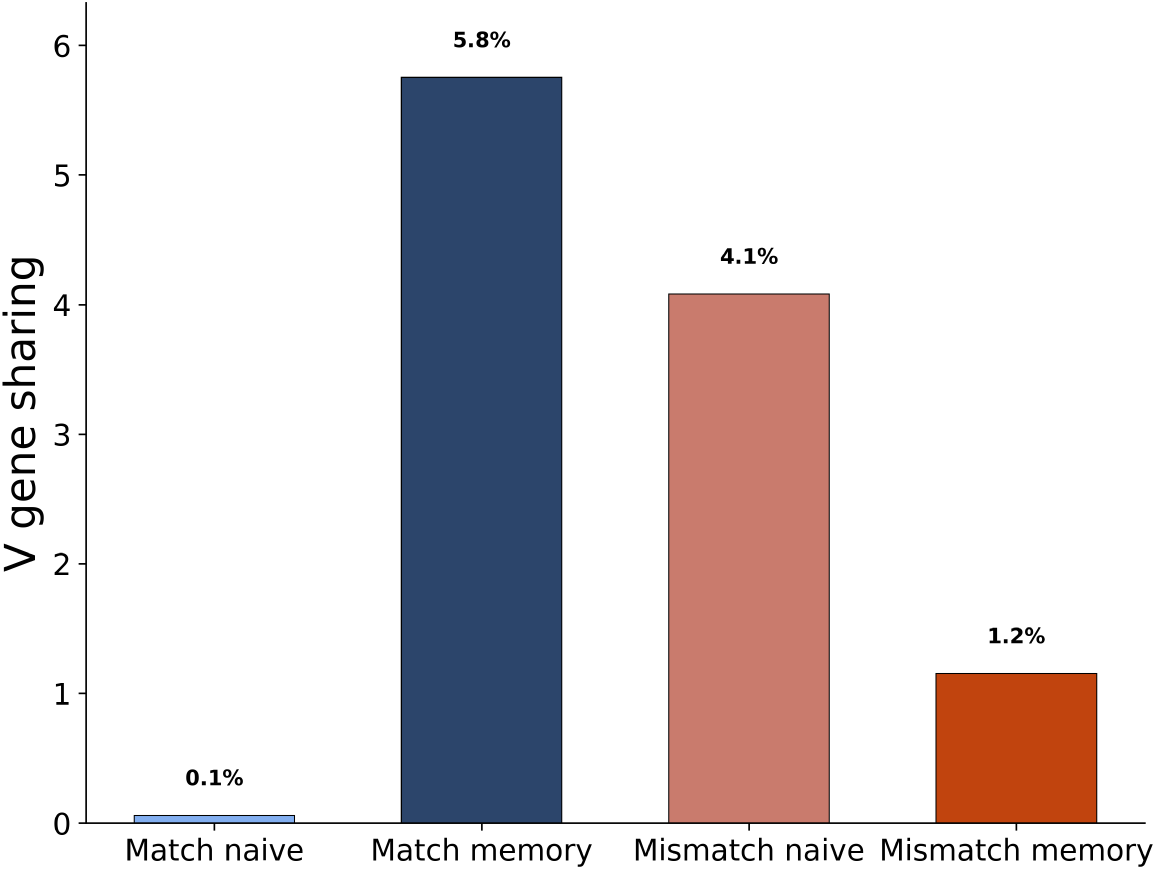
V gene constraint in conditionally generated LC sequences. Proportion of HCs for which *≥*80% of generated LCs utilize the same V gene. Higher percentages indicate more constrained V gene family usage during LC generation. Memory B cells demonstrate greater V gene constraint than naive B cells. Match naive: heavy and LCs both predicted as naive; Match memory: heavy and LCs both predicted as memory; Mismatch naive: naive HC paired with memory-predicted LC; Mismatch memory: memory HC paired with naive-predicted LC.

**Fig. A9.**
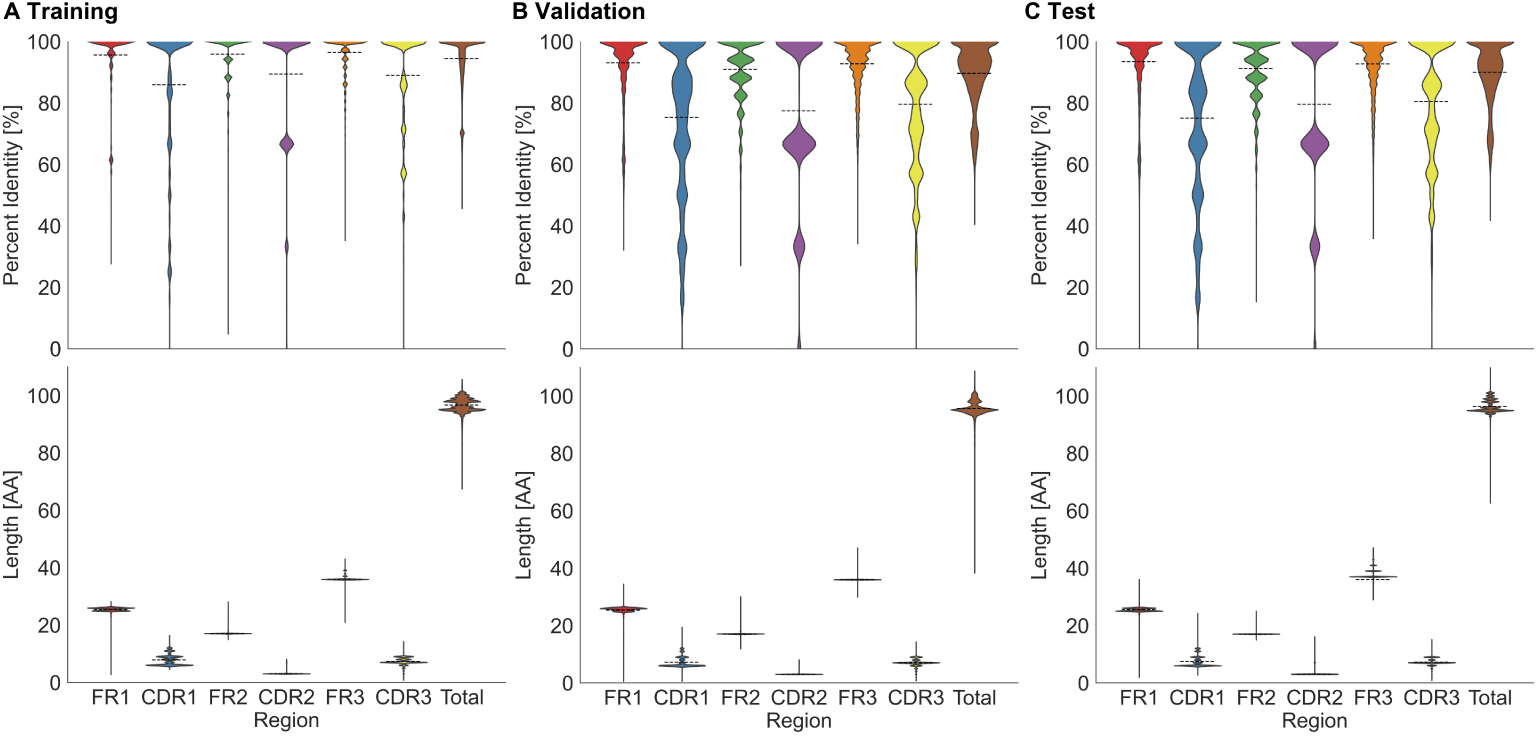
Distribution of germline identity and region lengths for LC sequences across dataset splits. For each dataset (A: Training set, B: Validation set, and C: Test set), violin plots show the distribution of mean percent identity to germline sequences (upper panel) and mean sequence lengths (lower panel) for framework regions (FR1-3) and complementarity-determining regions (CDR1-3). Black dashed lines indicate the mean values for each region.

**Fig. A10.**
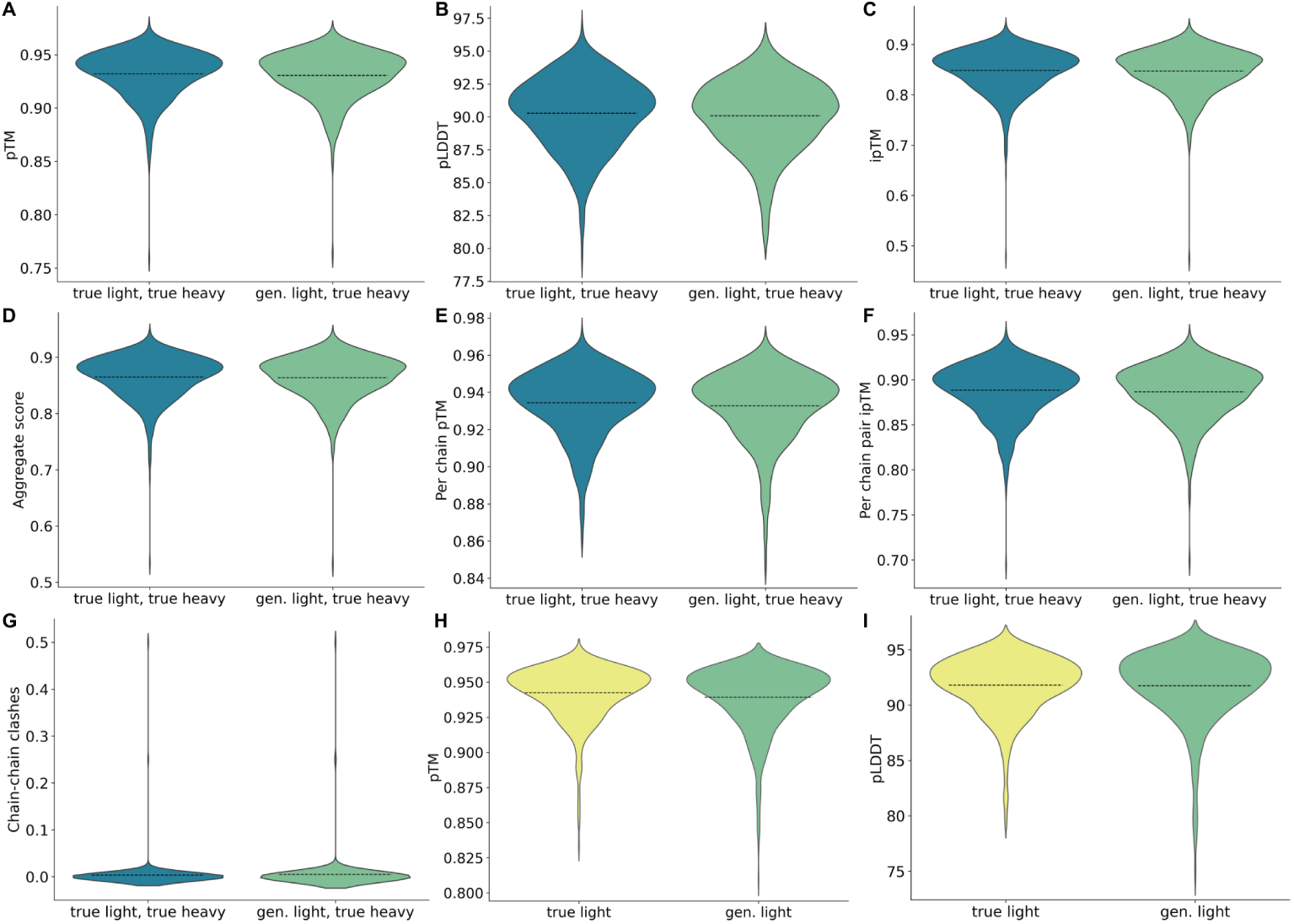
Structural assessment of HC-LC co-folding and individual LC folding using Chai-1 predictions. (A-G) Comparison of HC-LC co-folding quality between true HC-LC chain pairs (blue) and HCs co-folded with conditionally generated LCs (green) across multiple structural metrics. (H, I) Individual LC folding quality metrics for individual LC sequences, comparing true LCs (yellow) with conditionally generated LCs (green) folded independently.

**Table A9.** Structural quality assessment of true and generated LC sequences using Chai-1 predictions. Comparison of folding quality metrics for HC-LC co-folding (true LC-HC pairs vs. generated LC paired with true HC) and individual LC folding. Values represent mean scores across all evaluated sequences. Co-folding metrics include interface-specific measures (ipTM, per-chain pair ipTM) and chain clash analysis, while individual LC metrics focus on single-chain folding quality.

| Metric | Co-folding |  | Individual Light Chain |  |
| --- | --- | --- | --- | --- |
|  | True LC-HC | Gen. LC-HC | True LC | Gen. LC |
| Sample size | 731 | 568 | 509 | 682 |
| Per-chain pTM | 0.934 | 0.933 | — | — |
| Aggregate score | 0.865 | 0.864 | — | — |
| pTM | 0.932 | 0.931 | 0.942 | 0.939 |
| ipTM | 0.848 | 0.847 | — | — |
| Per-chain pair ipTM | 0.888 | 0.887 | — | — |
| Mean pLDDT | 90.27 | 90.08 | 91.79 | 91.74 |
| Chain-chain clashes | 0.003 | 0.005 | — | — |

**Table A10.** Hyperparameters and performance metrics of the Heavy2Light Encoder-Decoder model. . The model combines the HeavyBERTa encoder for HC representation learning with the LightGPT decoder for LC sequence generation.

| Training Configuration |  |
| --- | --- |
| Number of epochs | 50 |
| Batch size | 64 |
| Learning rate | $1e^{-5}$ |
| Weight decay | 0.1 |
| Optimizer | AdamW |
| Learning-rate scheduler | Linear |
| Gradient clipping (max-norm) | 1.0 |
| Model Architecture |  |
| Model type | Encoder-Decoder |
| Encoder | HeavyBERTa (RoBERTa-based) |
| Decoder | LightGPT (GPT-2-based) |
| Cross-attention | Enabled |
| Vocabulary size | 25 |
| *Encoder (HeavyBERTa)* |  |
| Hidden size | 512 |
| Layers | 4 |
| Attention heads | 4 |
| Intermediate size | 2048 |
| Max sequence length | 512 |
| *Decoder (LightGPT)* |  |
| Hidden size | 768 |
| Layers | 12 |
| Attention heads | 12 |
| Context length | 1024 |
| Generation Parameters |  |
| Sampling method | Nucleus (top-$p$) |
| Top-$p$ | 0.85 |
| Temperature | 0.8 |
| Top-$k$ | 0 |
| Max new tokens | 115 |
| Adapter Configuration |  |
| Adapter type | Bottleneck |
| Multi-head adapter | True |
| Output adapter | True |
| Reduction factor | 16 |
| Activation function | ReLU |
| Final Performance |  |
| Final training loss | 0.2078 |
| Final evaluation loss | 0.3828 |
| Total training steps | 367 750 |
| Training runtime | ~34.9 h |

**Table A11.** Hyperparameters and performance metrics for HeavyBERTa model configurations.

| Hyper-parameter / Metric | Small config. | Large config. |
| --- | --- | --- |
| *Training setup* |  |  |
| Number of epochs | 22 | 9.18 |
| Batch size | 16 | 16 |
| Learning rate | $5 \times 10^{-5}$ | $5 \times 10^{-5}$ |
| Weight decay | 0.1 | 0.1 |
| Optimizer | AdamW | AdamW |
| LR scheduler | Linear | Linear |
| *Model architecture* |  |  |
| Model type | RobertaForMaskedLM |  |
| Vocabulary size | 25 | 25 |
| Max sequence length | 512 | 512 |
| Max position embeddings | 514 | 514 |
| Position embedding type | Absolute | Absolute |
| Hidden activation | GELU | GELU |
| Hidden size | 512 | 768 |
| Layers | 4 | 12 |
| Attention heads | 4 | 12 |
| Intermediate size | 2048 | 3072 |
| *Performance* |  |  |
| Final training loss | 0.4476 | 0.4379 |
| Final evaluation loss | 0.4376 | 0.4234 |
| Evaluation accuracy | 0.8895 | 0.8912 |
| Total training steps | 136 254 500 | 56 832 000 |
| Training runtime | ~244 h | ~673 h |

**Table A12.** Hyperparameters and performance metrics for the LightGPT model.

| Training Configuration |  |
| --- | --- |
| Number of epochs | 41 |
| Batch size / total batch size | 16 / 32 |
| Learning rate | $5 \times 10^{-5}$ |
| Weight decay | 0.1 |
| Optimizer | AdamW |
| LR scheduler | Linear |
| Model Architecture |  |
| Model type | GPT2LMHeadModel |
| Vocabulary size | 25 |
| Context length / max positions | 1024 |
| Hidden activation | `gelu_new` |
| Hidden size | 768 |
| Layers | 12 |
| Attention heads | 12 |
| Inner dimension | 3072 |
| Embedding / attention / residual dropout | 0.1 / 0.1 / 0.1 |
| Layer-norm $\epsilon$ | $1 \times 10^{-5}$ |
| Initializer range | 0.02 |
| Performance |  |
| Final training loss | 0.3517 |
| Final evaluation loss | 0.5446 |
| Evaluation accuracy | 0.8635 |
| Total training steps | 12 265 000 |
| Training runtime | ~606 h |

**Table A13.** Training parameters for naive/memory B cell classification models. Hyperparameters used for fine-tuning HeavyBERTa to classify HC sequences and LightGPT to classify LC sequences by B cell developmental state (naive vs. memory).

| Parameter | HeavyBERTa classifier | LightGPT classifier |
| --- | --- | --- |
| Max length | 150 | 150 |
| Batch size | 64 | 64 |
| Epochs | 200 | 50 |
| Learning rate | $3e^{-6}$ | $3e^{-6}$ |
| Weight decay | 0.01 | 0.01 |
| Dropout | 0.1 | 0.3 |

